# Phylogenomics and species delimitation in the brown snake *Storeria dekayi* (Natricidae)

**DOI:** 10.1101/2025.10.30.685713

**Authors:** Victor Gabriel Castillo-Sanchez, Adrian Nieto-Montes de Oca, John J. Wiens, Marysol Trujano-Ortega, Uri O. García-Vazquez

**Author notes:** Corresponding author at: Laboratorio de Sistematica Molecular, Carrera de Biología, Unidad de Investigación Experimental Zaragoza, Facultad de Estudios Superiores Zaragoza, Universidad Nacional Autónoma de Mexico, Batalla 5 de Mayo s/n, Col. Ejercito de Oriente, 09230, CDMX, Mexico.

## Abstract

The brown snake, *Storeria dekayi*, is distributed across southeastern Canada, the eastern United States, and eastern Mexico, with isolated records in Central America. Despite its broad range, *S. dekayi* is currently recognized as a single species. Historically, eight subspecies were described based primarily on head coloration patterns, but subsequent genomic analyses of populations from the U.S.A led to their synonymization. Recently, a population discovered in the Cuatro Cienegas Basin, Coahuila, Mexico has been suggested to represent an undescribed endemic species. The aim of this study is to investigate the relationships within *S. dekayi*, with an emphasis on the previously unstudied populations from Mexico and Central America. To this end, we generated genomic data (ddRADseq) to conduct phylogenetic analyses, species-tree estimation, population-structure analyses, and species delimitation. Concatenated likelihood analyses estimated a tree in which populations of *S. dekayi* from the south-central U.S. were paraphyletic relative to an undescribed species from the Cuatro Cienegas Basin. Population structure and species delimitation analyses suggest the existence of three distinct species, including the populations from eastern North America (recognized as *S. dekayi*), the population from Cuatro Cienegas, and the populations of *S. dekayi* from eastern Mexico and Central America. However, we conservatively recognize only two species.

## 1. Introduction

The discovery of new species is essential for biodiversity conservation and understanding the speciation process (Yi et al., 2023). Cryptic species are defined as genetically distinct lineages that are morphologically similar (Bickford et al., 2007; Hending, 2025). However, substantial variation in coloration patterns may occur without corresponding genetic divergence (Lifjeld, 2015; Pyron et al., 2016), potentially leading to misclassifications (Cox and Davis Rabosky, 2013). The integration of molecular data in species delimitation has been crucial for uncovering cryptic species that are widely distributed across diverse taxonomic groups (Nygren, 2014; Perez-Ponce de León and Poulin, 2016; Fišer et al., 2018; Li and Wiens, 2023; Hending, 2025).

The discovery of cryptic species can be challenging when there is limited resolution of the tree based on certain molecular markers (Quattrini et al., 2019). For instance, incomplete lineage sorting and historical hybridization can complicate species delimitation when relying on mitochondrial DNA data as the only type of evidence, particularly in recently diverged taxa (Hickerson et al., 2006; Zheng et al., 2017; Steenwyk et al., 2023). Consequently, genome-wide data provide a powerful tool to detect gene-tree discordance and investigate the underlying causes of phylogenetic incongruence during rapid diversification events (Arcila et al., 2017).

Advances in genomic-scale DNA sequencing technologies have significantly increased the data available to help infer species boundaries (Parker et al., 2022). Among these, Double Digest Restriction-Site Associated DNA Sequencing (ddRADseq) is a next-generation sequencing approach that enables the analysis of phylogeny, species limits, and population genetic structure by detecting thousands of single nucleotide polymorphisms (SNPs) across the genome (Rowe et al., 2011; Peterson et al., 2012; Wang et al., 2022). These data have been instrumental in helping to resolve species complexes using phylogenetic and species-delimitation methods (Nieto-Montes de Oca et al., 2017; Barley et al., 2024; Burriel-Carranza et al., 2025). Furthermore, genomic data provide substantial power for detecting genetic structure within species, thereby enhancing our understanding of their genetic diversity (Lavretsky et al., 2019; Piñeros et al., 2022).

The brown snake (*Storeria dekayi*; Holbrook, 1839) has a broad distribution from Canada to Honduras. In eastern North America, it occurs from southern Ontario and Quebec in Canada and south to southern Florida, and west to eastern Texas in the United States. In Mexico, it occurs in the eastern part of the country, spanning from the northern states of Nuevo León and Tamaulipas to southern states including Veracruz, Puebla, and Chiapas. Additionally, isolated records have been reported from Central America, specifically in southern Guatemala and central Honduras (Wallach et al., 2014; Uetz and Hosek, 2022).

Eight subspecies of *Storeria dekayi* have been described across this range (Fig. 1). These include (type localities in parentheses), *S. d. anomala* Dugès, 1888 (Orizaba, Veracruz, Mexico); *S. d. dekayi* (Massachusetts, U.S.A.); *S. d. limnetes* Anderson, 1961 (St. Charles Parish, Louisiana, U.S.A.); *S. d. temporalineata* Trapido, 1944 (San Rafael, Jicaltepec, Veracruz, Mexico); *S. d. texana* Trapido, 1944 (Kendall County, Texas, U.S.A.); *S. d. tropica* Cope, 1885 (Peten, Guatemala); *S. d. victa* Hay, 1892 (Florida, U.S.A.); and *S. d. wrightorum* Trapido, 1944 (Reelfoot Lake, Tennessee, U.S.A.). These subspecies were originally distinguished primarily based on coloration patterns and scale counts, with a general trend of higher ventral scale counts in more southern subspecies. Recent phylogenomic and morphological analyses have led to the synonymization of several of these taxa, with the exception of *S. d. victa*, which was elevated to a distinct species (Pyron et al., 2016). However, no populations from Mexico and Central America were included in these analyses, making it unclear whether the subspecies occurring there should be synonymized or instead recognized as distinct species.

**Figure 1.**
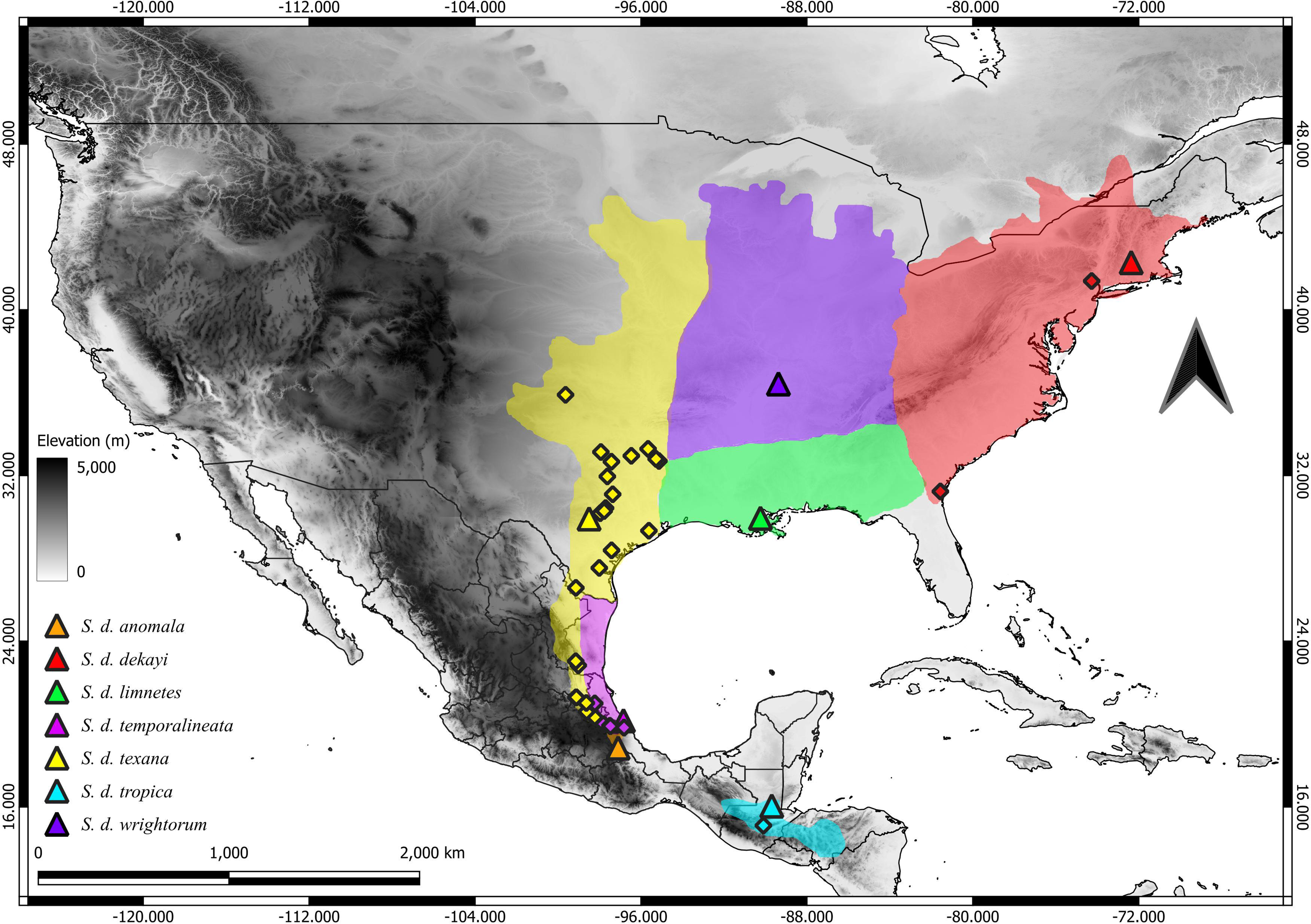
Type localities of *Storeria dekayi* subspecies (triangles) and the genetic samples used in this study (diamonds). The approximate geographic distributions of *S*. *dekayi* subspecies (Trapido, 1944) are shaded to illustrate geographic coverage of sampling.

Recently, García-Vazquez et al. (2019) reported the presence of an isolated *Storeria* population in the Cuatro Cienegas Basin (CCB) in Coahuila, Mexico, exhibiting a scale count pattern similar to that of *S. dekayi*. Based on phylogenetic analyses of mitochondrial genes (ND2) and morphological differentiation, Vite-Hernandez (2023) concluded that this population represents a distinct lineage that is closely related to *S. dekayi*. This species has unique morphological characteristics relative to other *Storeria*, including a gray dorsum, a pale venter lacking dark spots, and a smaller body size. Consequently, he proposed recognizing this population as a new *Storeria* species, that was likely endemic to the CCB. This pattern of endemism has been observed in other reptile species, including *Scincella kikaapoa* García-Vazquez et al. 2010, *Gerrhonotus mccoyi* García-Vazquez et al. 2018, and *Terrapene coahuila* Schmidt and Owens, 1944.

This study aims to re-evaluate the species boundaries of *Storeria dekayi* populations, specifically those from Mexico and Central America. Previous studies addressing the taxonomy of this group relied on different types of molecular data and limited sampling, leading to inconclusive results regarding species limits. For example, Hernandez-Vite (2023) analyzed mitochondrial DNA sequences, whereas Pyron et al. (2016) used anchored hybrid enrichment (AHE) loci from North American populations. To address this gap, we use ddRADseq data from extensive sampling of *S*. *dekayi* populations and other *Storeria* species, to infer their phylogenetic relationships and species limits. Our working hypothesis is that Mexican and Central American populations may represent distinct evolutionary lineages, potentially warranting taxonomic recognition.

## 2. Methods

### 2.1. Taxon sampling

To analyze the southernmost populations of *Storeria dekayi*, we focused our RADseq sampling on southern populations not included by Pyron et al. (2016). We sampled 45 individuals from the genus (Table 1; Fig. 1), including 38 specimens of *S. dekayi* sensu Pyron et al. (2016) representing the historical subspecies *S. d. dekayi* (*n*=2 individuals)*, S. d. texana* (*n*=29)*, S. d. temporalineata* (*n*=6), and *S. d. tropica* (*n*=1). Additionally, we included five individuals of *S. occipitomaculata*, four of *S. storerioides*, one of *S. victa* and two of *Storeria* sp. CCB (Table 1). To root the phylogenetic tree, we used one sample of *Thamnophis scalaris* as an outgroup (Nuñez et al., 2023).

**Table 1.**
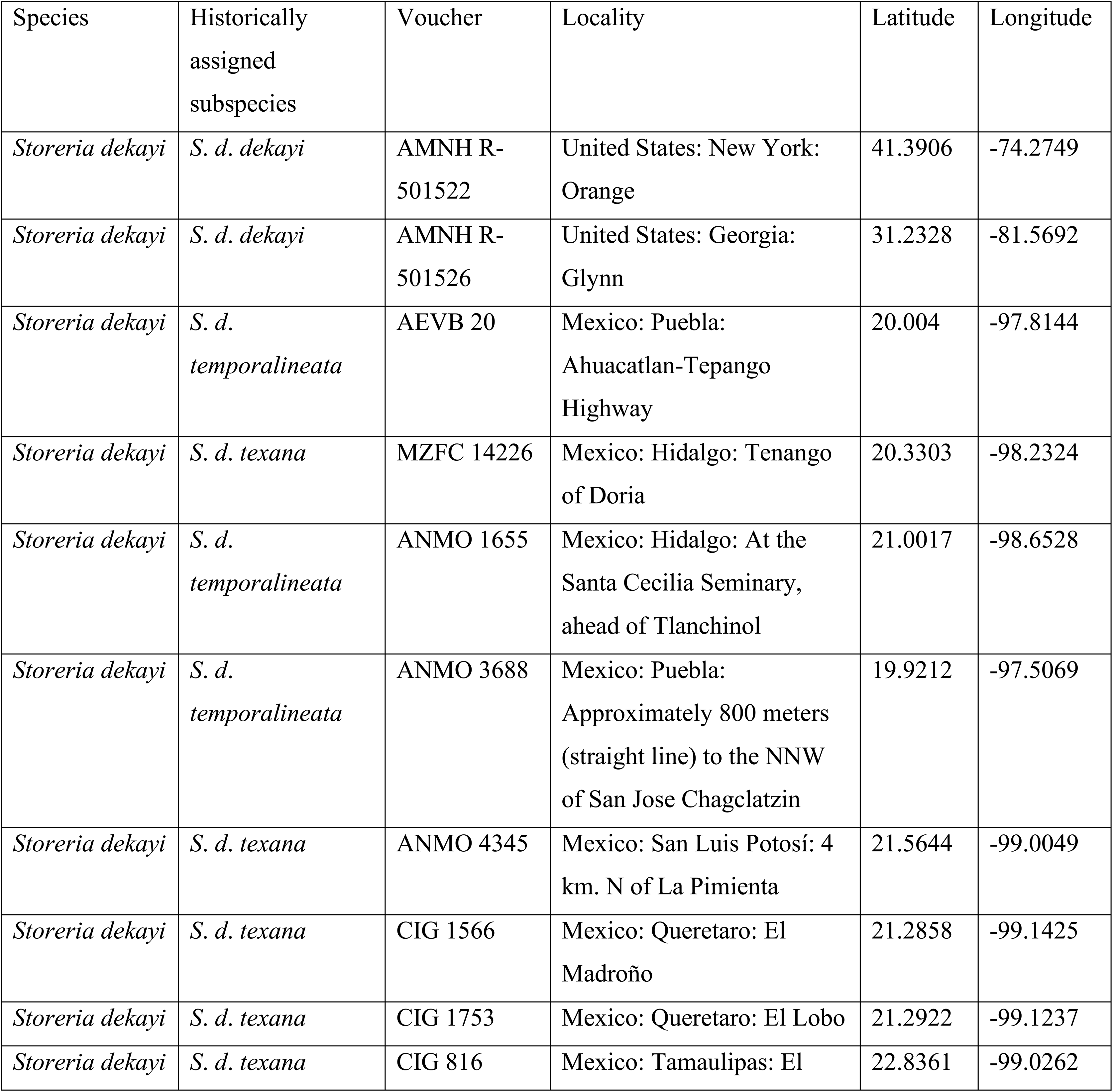
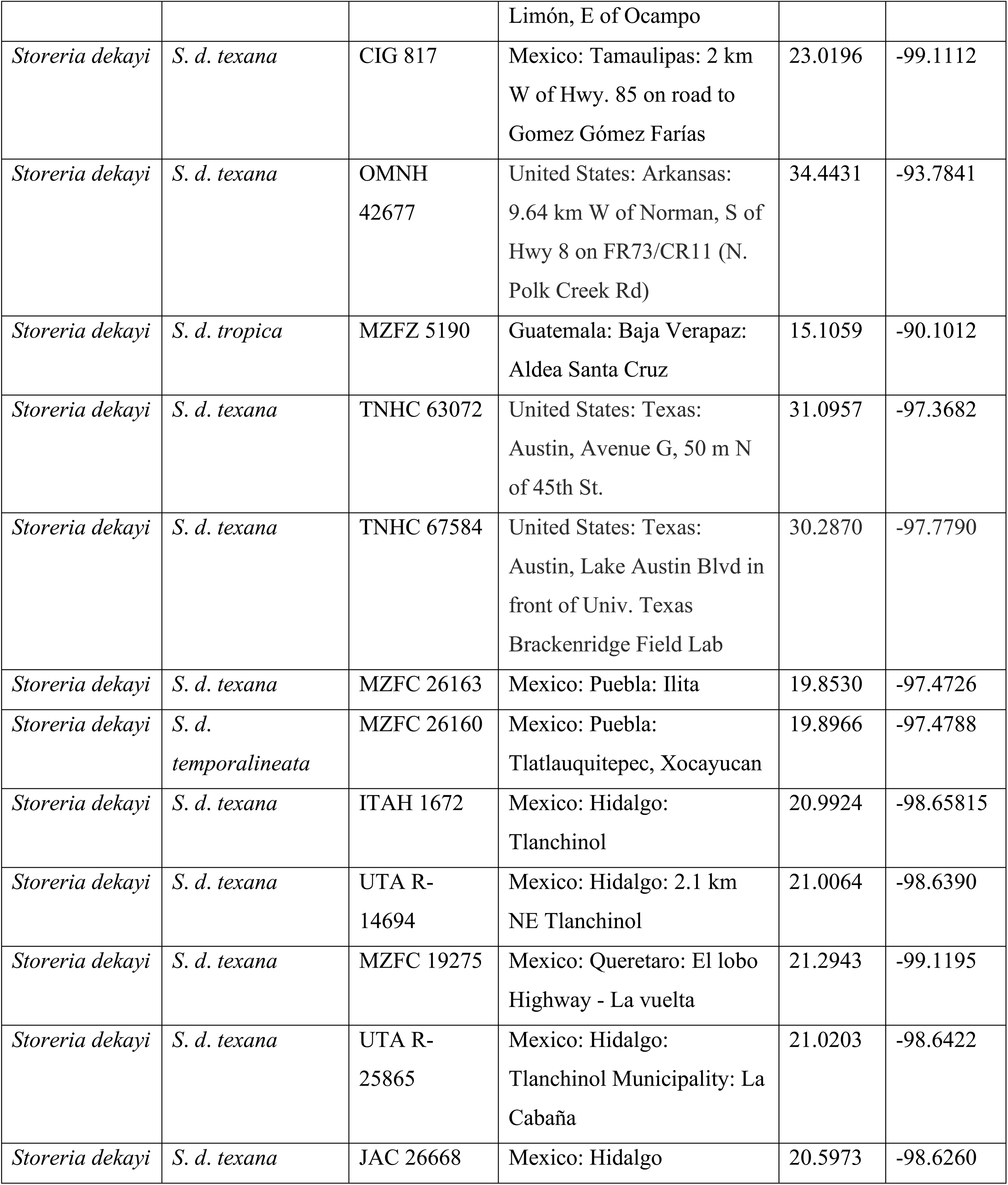
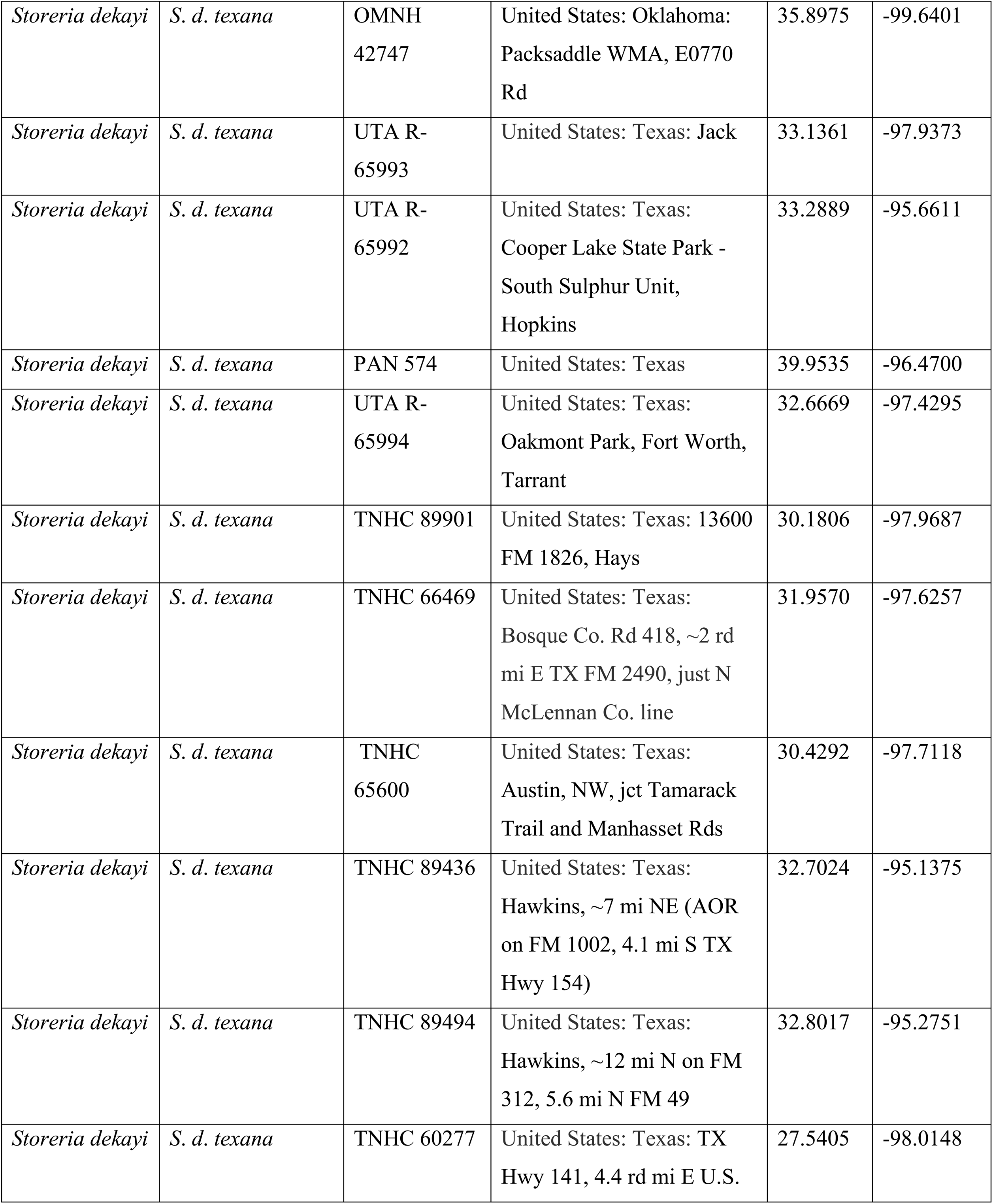
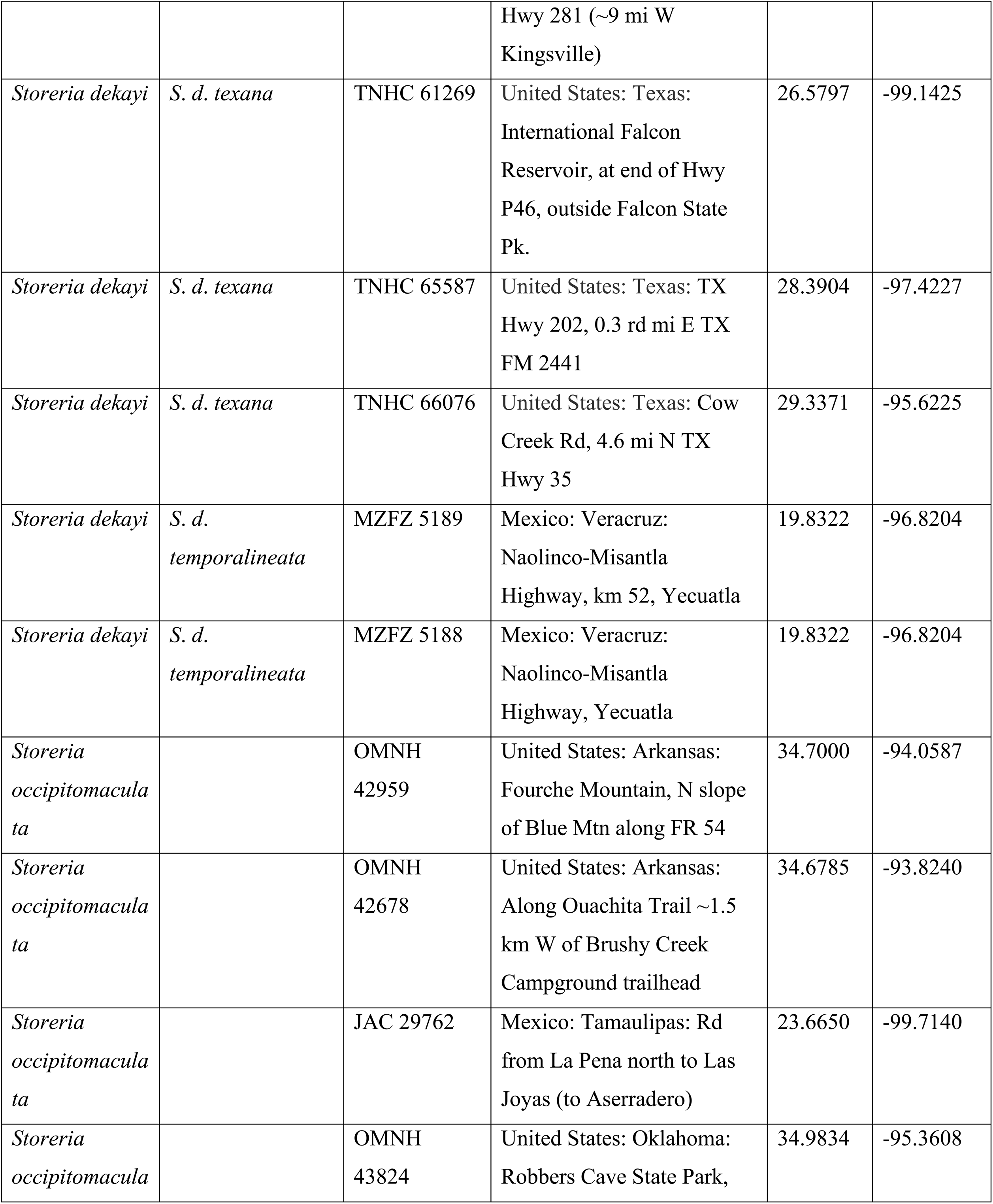
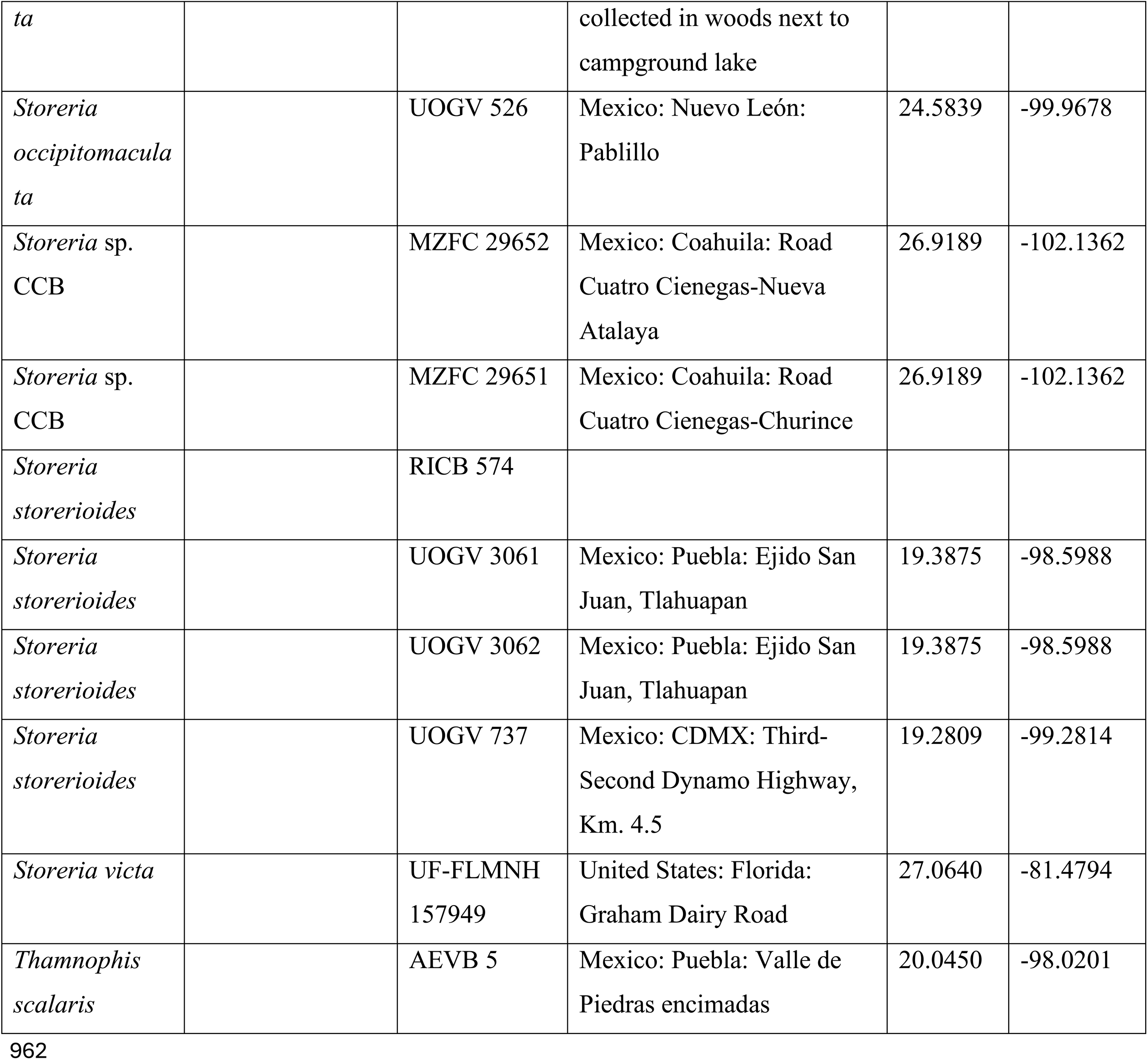
Taxon sampling for ddRAD sequence data collection. Institutional abbreviations for museums and collections follow Sabaj (2023), except for MZFZ (Museo de Zoología of the Facultad de Estudios Superiores Zaragoza, Universidad Nacional Autónoma de Mexico); ANMO, AEVB, JAC and CIG numbers are field numbers for specimens to be catalogued in the MZFZ.

### 2.2. Molecular procedures

#### 2.2.1. DNA extraction

Whole genomic DNA was extracted from tissue samples using the Qiagen DNeasy Kit (Qiagen Inc.). Prior to library preparation, DNA concentration was quantified using a Qubit fluorometer (Thermo Fisher Scientific) and quality was visually assessed on a 1% agarose gel.

#### 2.2.2. ddRADseq libraries

We generated ddRADseq libraries using a modified version of the protocol from Peterson et al. (2012). Genomic DNA from each sample was digested using the restriction enzymes SbfI (recognition site 5′–CCTGCAGG–3′) and MspI (recognition site 5′–CCGG–3′). The digestion was performed by incubating the samples at 37°C for two hours in a thermocycler. The reaction mixture consisted of 26 μl of DNA (250 ng), 2.5 μl of 10× Cutsmart® buffer, 0.25 μl of MspI (20 U/μl), and 0.5 μl of SbfI (20 U/μl). The digestion products were then purified using 1.5× Sera-Mag Magnetic SpeedBeads.

We then ligated the DNA fragments to specific adapters for SbfI and MspI. The ligation reaction was performed in a solution containing 30 μl of digested DNA, 1.5 μl of each specific adapter, 3.4 μl of 10× Ligase Buffer (NEB®), and 0.6 μl of T4 DNA Ligase (400 U/μl). This reaction was incubated at 22°C for 40 minutes, followed by 10 minutes at 65°C. The ligated products were purified using 1.5× Sera-Mag Magnetic SpeedBeads.

We then amplified the restriction-ligation products using the polymerase chain reaction (PCR) with primers specific for the Illumina sequencer. Each PCR reaction contained 20 μl of ligated DNA, 1.25 μM of specific adapters, 0.6 mM dNTPs, 6 μl of 5× Phusion Buffer HF, 0.3 μl of Phusion DNA Polymerase, and 0.6 μl of nuclease-free water. PCR conditions consisted of an initial denaturation at 94°C for 3 minutes, followed by 15 cycles of 94°C for 25 seconds, 57°C for 20 seconds, and 72°C for 30 seconds, with a final extension at 72°C for 10 minutes.

Finally, the ddRADseq libraries were purified using 1.5× Sera-Mag Magnetic SpeedBeads. Library construction success was assessed using a Qubit Fluorometer, and DNA size selection (470–630 bp) was performed using a Pippin Prep automated size selector (Sage Science,®). The ddRADseq libraries were sequenced at Macrogen, Inc. using the Illumina HiSeq 2500 platform.

#### 2.2.3. Bioinformatic processing

We processed the ddRADseq data using the ipyrad pipeline (version 0.7.19; Eaton and Overcast, 2020). Reads were filtered using a quality-score threshold of 28. Illumina adapters were removed using the trimming option, and the resulting de novo clusters were aligned with Muscle version 3.8.31 (Edgar, 2004).

To estimate the optimal clustering threshold (CT) that minimizes false homozygosity and heterozygosity (i.e., the optimal CT), we generated assemblies in ipyrad whit default values, using eight different CT values (set at increments of 0.2 between 0.84 and 0.98) and then evaluated five metrics based on population and landscape genetics, as proposed by McCartney-Melstad et al. (2019). These five metrics were (Figs. S1 and S2): (1) percentage of loci inferred as paralogues; (2) per-individual percent heterozygosity; (3) total number of variable sites (single-nucleotide polymorphisms; SNPs); (4) cumulative variance explained by the first eight principal components of the genetic data (PCs); and (5) Pearson’s correlation coefficient between pairwise genetic dissimilarity and the percentage of missing data. Metrics 1 and 2 identified 94% as the optimal sequence similarity threshold for our dataset, while both Pearson’s correlation coefficient and the cumulative variance explained by the first eight PCs favored an 84% threshold. Metric 3, in turn, identified an optimal clustering threshold of 92%. Based on these results, and following the approach proposed by McCartney-Melstad et al. (2019), we clustered and aligned the data using a 92% threshold (Figs. S1 and S2). We set the minimum number of samples per locus to 40% for phylogenetic analysis (Fig. S3). The final dataset comprised 875,462 base pairs and 8,519 unlinked SNPs, with 42.04% missing data. Raw Illumina reads are available in the GenBank sequence read archive (SRA) under the BioProject accession XXX.

#### 2.2.4. Population structure

For analyses that did not require an outgroup, we performed a de novo assembly of the *Storeria dekayi* samples. Excluding the outgroup increases the number of shared loci; therefore, we generated additional datasets using different clustering thresholds (Table S1, Supplementary Data), following McCartney-Melstad et al. (2019). The resulting Variant Call Format (VCF) file was filtered using VCFtools version 0.1.17 (Danecek et al., 2011) to retain only unlinked, biallelic SNPs under the following criteria: --min-alleles 2, --max-alleles 2, --maf 0.1, and --max-missing 0.5. The filtered VCF file was then converted into a .str file for STRUCTURE analysis using the program PGDSpider version 2.1.1.5 (Lischer and Excoffier, 2012). A second filtering was performed without applying the --maf option, since the influence of minor allele frequency (MAF) can affect the assignment of individuals to an ancestral cluster (Linck and Battey, 2019). For this purpose, a file was generated using the --singletons option in VCFtools, which details the location of singletons across individuals. The filtering was then carried out with the criteria: - -min-alleles 2, --max-alleles 2, --exclude-positions, and --max-missing 0.5 to compare the efficiency of both filtering approaches.

We implemented two approaches to investigate population structure. The first was based on non-parametric statistics using principal component analysis (PCA) in R with the package *SNPrelate* (Zheng et al., 2012). Data visualization was performed with the R package *tidyverse* (version 1.3.0; Wickham et al., 2019).

The second approach employed a Bayesian clustering algorithm implemented in the program STRUCTURE version 2.3.4 (Pritchard et al., 2000) to determine the optimal number of genetic clusters (K). We ran 10 replicate analyses for each value of K, each with 300,000 generations and discarding the first 100,000 generations as burn-in. The optimal number of clusters (K) was estimated using the method of Evanno et al. (2005), as implemented in the program Structure Harvester (Earl and Vonholsdt, 2012), which determines the optimal K based on the lnPr(D|K) statistic (Pritchard et al., 2000). Results from the replicate runs were summarized and visualized using CLUMPP version 1.1.2 (Jakobsson and Rosenberg, 2007) and RStudio version 2024.12.1 (R Core Team, 2021).

#### 2.2.5. Phylogenetic analyses

Using the identified clustering threshold, we generated a series of assemblies by adjusting the minimum number of samples per locus parameter in ipyrad (Eaton and Overcast, 2020). This parameter defines the minimum number of samples that must have data at a given locus for it to be retained in the final dataset (Table S2). Different data matrices were generated at the selected threshold, evaluating values from 30% to 80% of total number of taxa (in 10% increments) for the minimum number of samples per locus parameter (Lara-Tufiño et al., 2025).

We performed concatenated maximum likelihood (ML) analyses of the generated datasets using the software RAxML-NG v.1.2.0 (Kozlov et al., 2019). The concatenated datasets included all loci, including both SNPs and invariant sites to improve branch length estimation and topological precision (Leache et al., 2015). We employed a GTR+Γ model and 100 starting trees (50 random and 50 parsimony) and assessed node support via a non-parametric bootstrap of 100 replicates, executed through CIPRES Sci., Gateway version 3.3 (Miller et al., 2010). Then, to select the optimal tree, the average bootstrap (BS) values for each resulting tree were calculated using the R package *ape* version 5.7-1 (Paradis and Schliep, 2019) and visualized with the function “boxplot()” in RStudio version 2024.12.1 interface (RStudio Team, 2020). These results are shown in Fig. S3 (Supplementary Data). Phylogenetic trees were visualized and edited using FigTree version 1.4.4 (Rambaut, 2018) and Inkscape version 1.2.

#### 2.2.6. Species tree analyses

The STRUCTURE analyses revealed that five samples from Arkansas, Oklahoma, and Texas (PAN 540, SMOMNH 42677, SMOMNH 42747, and TJL 2304–2305) likely exhibited introgression from the eastern and south-central U.S.A. populations, preventing their assignment to a single ancestral population. Consequently, we performed all subsequent analyses excluding samples with less than 60% assignment to any population, as we suspect that these individuals originated from each of the genetic clusters (Barley et al., 2024; Pang and Zhang, 2024).

Species trees for genetically distinct populations of *Storeria dekayi* were inferred using two methods, with samples assigned to putative species based on the results from the ML, STRUCTURE, and PCA analyses. First, we inferred a species tree topology using SVDquartets, implemented in PAUP version 4.0a (Swofford, 2001; Chifman and Kubatko, 2014), based on the “.usnps” output file generated by ipyrad, which contains unlinked SNPs. All possible quartets were evaluated, and node support was assessed via bootstrapping with 1,000 replicates. The sample of *S. victa* was excluded due to poor sequence quality and fewer loci recovered compared to other samples.

Next, we estimated a species tree using Bayesian inference in BPP version 4.7 (Yang, 2015; Yang and Rannala, 2010, 2014). We performed two independent species-tree analyses (A01 analysis in BPP), each starting from a different initial topology. Because a larger number of loci showed poor admixture and failed to reach convergence (following Barley et al., 2024), a dataset comprising 500 loci was used. This reduced set of loci also helped balance computational demands. According to the results of the STRUCTURE analysis, we excluded individuals with less than 60% of their genetic ancestry derived from a single genetic group to avoid phylogenetic uncertainty arising from the inclusion of potential hybrids (Pavón-Vazquez et al., 2024). Analyses used the “.alleles” file of ipyrad. Population size parameters (θ) were assigned an inverse-gamma prior IG (3, 0.0012), based on the mean uncorrected pairwise genetic distance (p-distance) within taxa. The divergence time at the root of the species tree (t₀) was assigned an inverse-gamma prior IG (3, 0.0303), based on the mean p-distance between the basal-most taxon (*S. occipitomaculata*) and the other taxa. Each analysis was set to take 200,000 samples with a burn-in of 20,000 samples and, a sampling frequency of 10.

#### 2.2.7. Isolation by distance and species delimitation

We compared geographic and genetic distances to test for evidence of isolation-by-distance (IBD) using SNP data and the R package *BEDASSLE* (Bradburd et al., 2013). To calculate a matrix of geographic distances, we used the R package *geosphere* (Hijmans et al., 2024). For this analysis, samples were assigned to populations based on the results of the STRUCTURE analysis. When groups belong to the same species, then genetic and geographic distances typically exhibit a continuous relationship. In contrast, a discontinuous relationship between genetic and geographic distances, both within and between groups, can be indicative of the presence of distinct species (Prates et al., 2024).

We conducted two hierarchical heuristic species delimitation under the MSC-M model including and excluding the five admixed samples, using the program Hierarchical Heuristic Species Delimitations (hhsd) version 0.9.9 (Kornai et al., 2023) to automate the estimation of the genealogical divergence index (gdi) between populations in BPP (Jackson et al., 2017; Leache et al., 2019). The species tree topology estimated in the BPP A01 analysis was used as the guide tree for delimitation. Bidirectional migration was allowed between all sister-species pairs in the phylogeny, including historical migration between common ancestors and their sister lineages. The migration rate prior was modeled using a gamma distribution G (0.1, 10), with a mean of 0.01 migrant individuals per generation following Kornai et al. (2023).

To estimate species divergence times and population sizes under the multispecies coalescent model, we performed an A00 analysis in BPP version 4.7 (Yang, 2015; Flouri et al., 2018). This analysis was based on 500 loci and employed inverse gamma priors for θ (IG (3, 0.0012)) and τ₀ (IG (3, 0.0303)). The posterior means of the parameters obtained from the analysis were then used to calculate the gdi (Leache et al., 2019).

#### 2.2.8. Hybridization

To explore the possibility of gene flow as a cause of discordance between the ML and species trees, we used the PhyloNetworks program version 0.17 (Solís-Lemus et al., 2017) to infer phylogenetic networks under the multispecies network coalescent model (MSNC). A SNP assembly (one SNP per locus) was generated in ipyrad, incorporating representatives of *S. dekayi* and *S.* sp. from CCB, including samples that exhibited less than 60% total ancestry. Using this assembly, we created a table of quartet-level concordance factors (CF) with the function “SNPs2CF” in R (Olave and Meyer, 2020).

The concordance factors and the previously inferred species tree were then used as input for PhyloNetworks to search for phylogenetic networks, allowing a maximum number of reticulation events (hmax) ranging from 0 to 3 (Myers et al., 2022). For each search, 50 independent runs were conducted. To evaluate statistical support for the optimal network, defined as the network with the lowest negative log-likelihood score, we performed a bootstrap analysis with 100 replicates, each consisting of 20 runs. The functions tree Edges Bootstrap and hybrid Bootstrap Support were then used to map bootstrap proportions onto the edges and hybrid nodes of the best network (Lara-Tufiño et al., 2025).

#### 2.2.9. Divergence-time estimation

We used the BPP software to estimate divergence times in the *S. dekayi* complex. We estimated population sizes of contemporary and ancestral populations (θs) and divergence times (τs) on the fixed species tree (A01 analysis in BPP). The same individuals were used as for the species tree analyses, to control for potential gene flow between populations. We also used the same inverse gamma priors for θ and τ0. Analyses were based on 500 loci. We transformed the coalescent times (highest posterior densities [HPDs]) calculated by BPP to geological times (Mya) using the equation t= τ/r (Angelis and dos Reis, 2015; Dos Reis, 2016). In this equation, tau is the mean HPD of coalescent times and *r* is a neutral substitution rate, with a rate for natricids of 8.1x10^-10^ substitutions per site per year (Perry et al., 2018).

#### 2.2.10. Morphological comparisons

To investigate the morphological differentiation among the lineages identified within the complex using molecular data, we examined eight specimens from the CCB population, 11 specimens of *S*. *dekayi* from the U.S.A., and 65 specimens from Mexico and Central America. The list of specimens examined is provided in Table S3. The list of 18 morphological characters examined (16 morphometric, 2 meristic) is given in Table S4.

Scale nomenclature followed Trapido (1944). However, given the lack of variation in scalation among the reviewed specimens (variation was observed only in ventral and subcaudal scales), morphometric measurements of the scales were performed. These were observed using a stereomicroscope, and measurements were taken with a Lenfench digital caliper (0.01 mm precision) and recorded to the nearest 0.1 mm. Counts of ventral and subcaudal scales were also recorded (Table S4). To visualize morphological variation, we performed principal component analyses (PCA) on the data using the PCA function from the R package FactoMineR (Lê et al., 2008). We considered there to be evidence for morphologically distinct species if there was no overlap among putative in the PCA plots from a given pair of PCs, for one or more pairs of axes.

## 3. Results

### 3.1. Population structure

Filtering of the SNP dataset with the --maf option resulted in, 916 retained unlinked SNPs. The results obtained with STRUCTURE and SNPRelate are summarized in Figure 2. In the second filtering, where singletons were excluded, a matrix comprising 1,998 SNPs was obtained, and the corresponding PCA and STRUCTURE results are presented in Figure S4.

**Figure 2.**
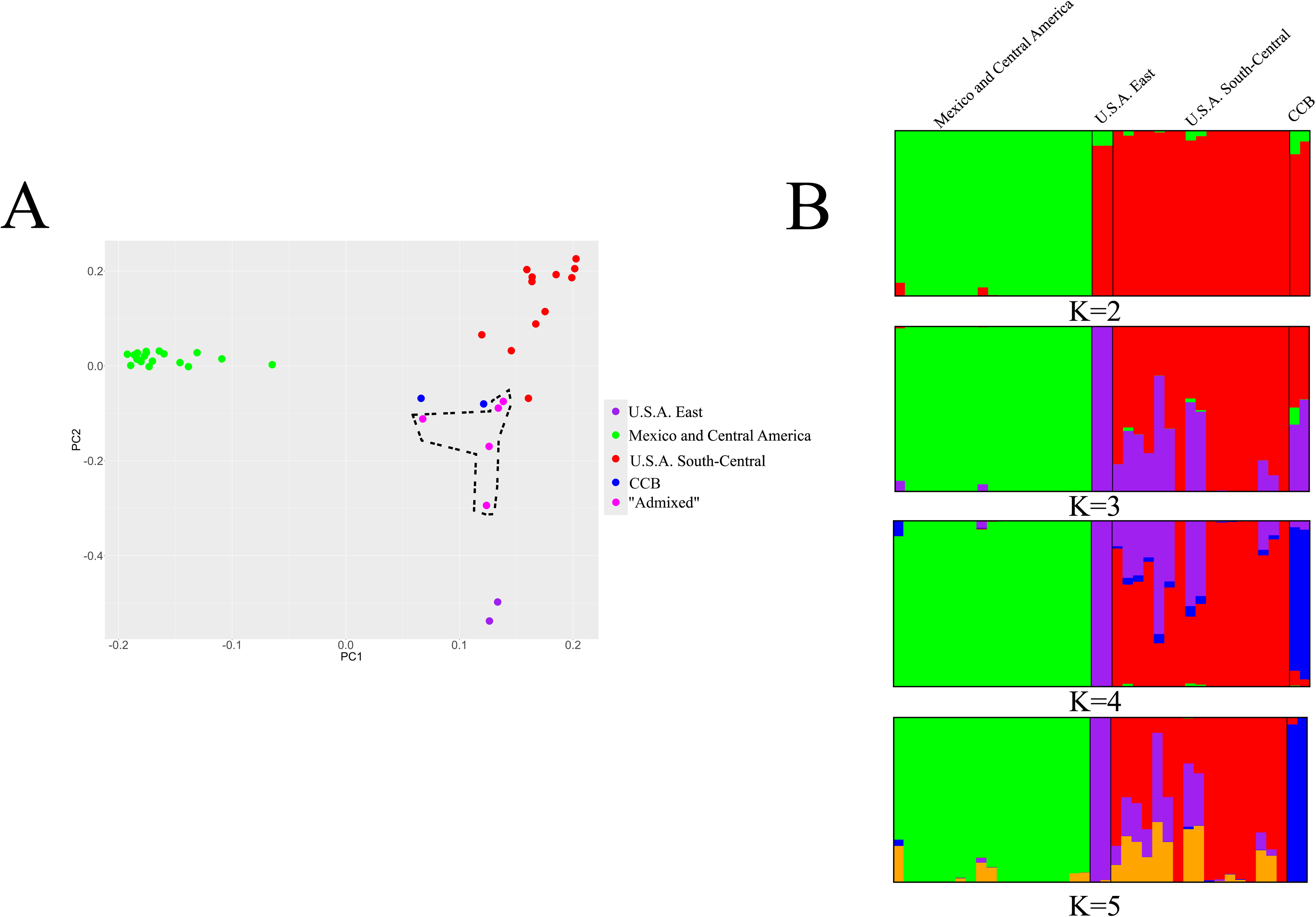
A. Population structure in *Storeria dekayi* inferred by means of Principal Component Analysis. Dots represent different individuals, colored by cluster. The five dots enclosed by the dashed line are highly admixed individuals from U.S.A. B. Population structure in *Storeria dekayi* inferred by STRUCTURE. Each vertical bar in the barplots represents an individual. K = 2 is the optimal K obtained with the delta K method, and K = 5 is the optimal K obtained with the ln Pr(D|k) method.

The analyses of both datasets yielded, similar patterns of structure among individuals. Using the ΔK method, the optimal value of K for the dataset filtered with the –maf option peaked at two (ΔK = 3,521.0559; Fig. S5A), whereas the dataset excluding singletons also yielded an optimal value of K of two (ΔK = 6,367.0741; Fig. S5C). In contrast, the ln Pr(D|K) method indicated an optimal value of K of five for both datasets (Figs. S5B, S5D). The five clusters consisted of the samples from: (1) Mexico and Central America, (2) CCB, (3) south-central U.S.A. (Arkansas, Texas, Oklahoma), (4) eastern U.S.A. (Georgia, New York), and (5) the samples displaying high admixture. The south-central U.S.A. group exhibited high admixture among different clusters at K = 3–5. Principal component analysis (PCA) of the RADseq data segregated the samples into three distinct clusters, corresponding to *S. dekayi* from: (1) Mexico and Central America, (2) eastern U.S.A., and (3) south-central U.S.A. The CCB haplotypes clustered with the highly admixed samples (Fig 2A). In the PCA, the first three PCs together explained 52.49% of the total variance, with PC1, PC2, and PC3 explaining 37.67%, 10.16%, and 4.66% of this variance, respectively.

### 3.2. Phylogenetic analysis

Our ML analysis of the concatenated data (Fig. 3) showed that *S. dekayi* and the CCB samples formed a strongly supported monophyletic group that the sister taxon to *S. victa*. Within the *S. dekayi* clade, the sampled populations from the eastern U.S.A. are sister taxon to all other *S. dekayi* populations plus the *S*. sp. CCB samples. The samples from south-central U.S.A., including five admixed individuals from south-central U.S.A., were paraphyletic with respect to the sister clades with the samples from Mexico and Central America and the CCB. Many of the relationships among the *S. dekayi* populations from south-central U.S.A. are strongly supported (BS >90%), including those showing the south-central U.S.A. populations to be paraphyletic.

**Figure 3.**
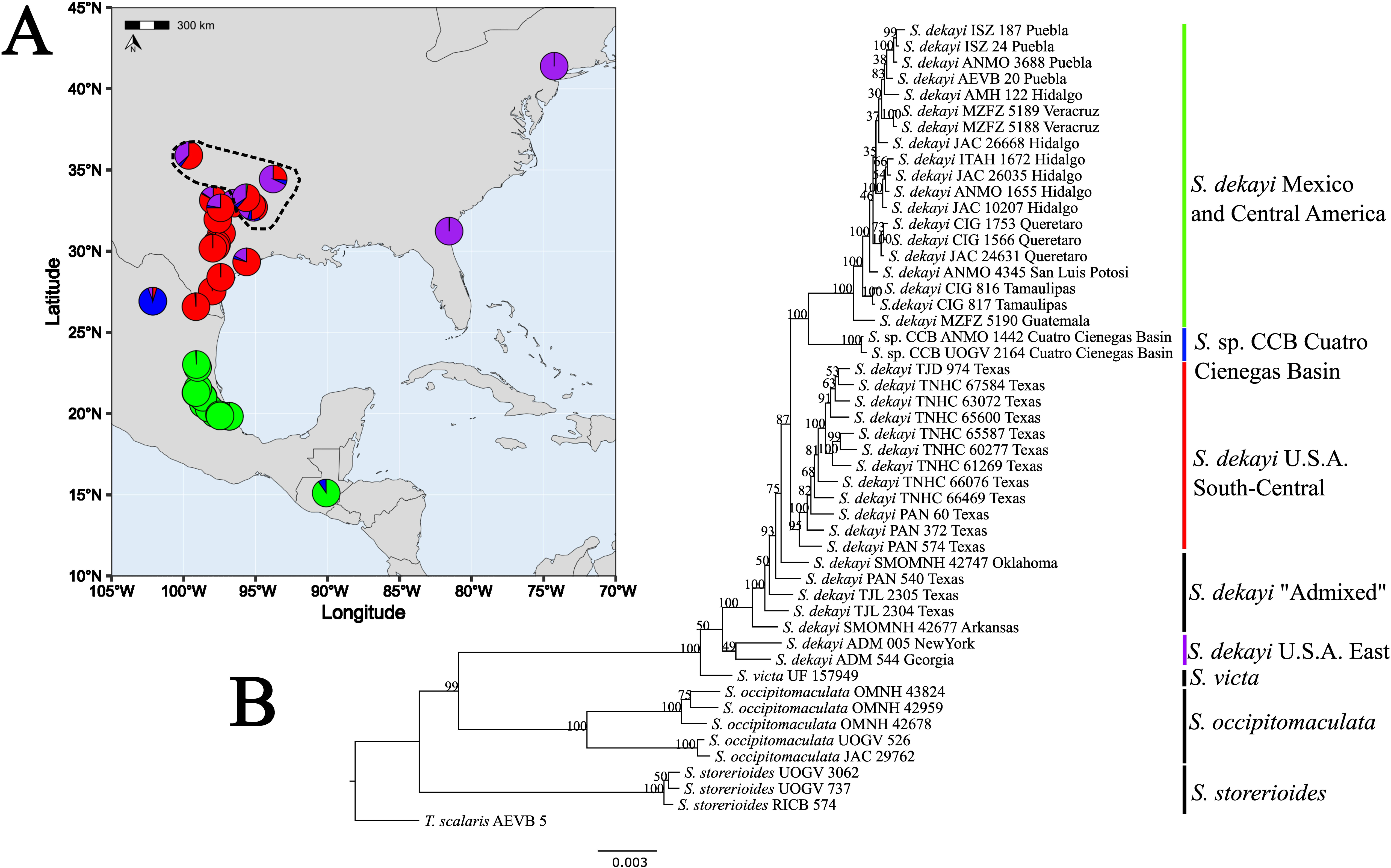
A. Geographic distribution of population structure in *Storeria dekayi* inferred with STRUCTURE. Each pie represents an individual, with multiple colors indicating mixed ancestry. Note the presence of five highly admixed individuals (“Admixed”) from U.S.A. enclosed by a dashed line. B. Phylogenetic tree for *Storeria* estimated with RAxML with a clustering threshold of 90% and minimum percentage of samples with data per locus = 40%. Numbers on nodes are bootstrap support (bs) values. Nodes with bs > 70 are considered well supported (Hillis and Bull, 1993).

### 3.3. Species tree analysis

In the species tree generated by SVDquartets (Fig. 4A), the five admixed samples were excluded, and the remaining individuals were assigned to putative species based on the clades recovered in the ML tree, the STRUCTURE results, and clusters in the PCA results. In the SVDquartets tree, *S. dekayi* from south-central U.S.A. formed the sister group to the lineage from the CCB, although this relationship was weakly supported (bootstrap support [BS] = 55). Together, these two groups formed the sister clade to *S. dekayi* from Mexico and Central America (BS = 100), and all of these taxa comprised the sister taxon to *S. dekayi* from eastern U.S.A. (BS = 100). However, we recognize that it is potentially problematic to treat the samples from south-central U.S.A. as a clade.

**Figure 4.**
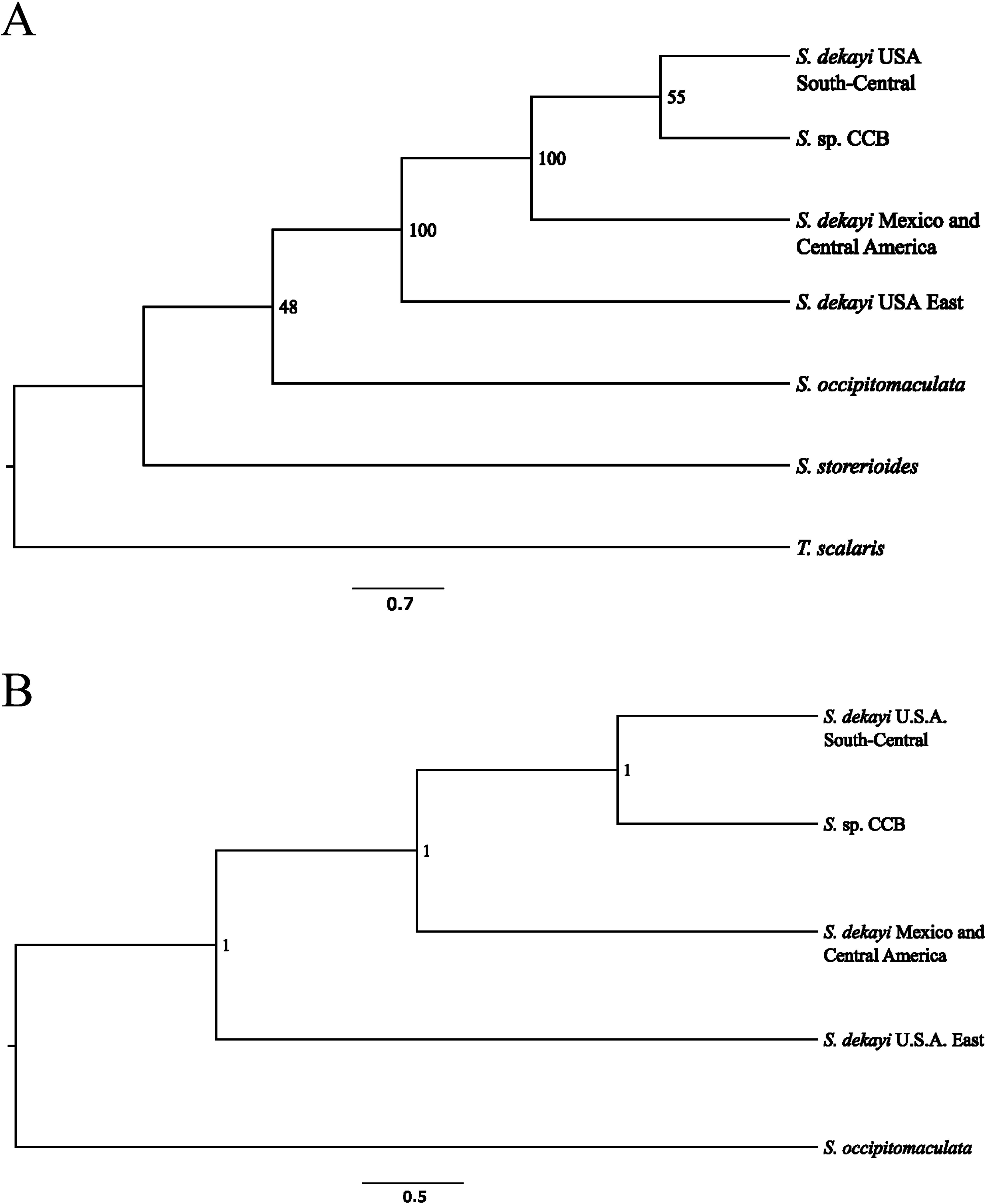
Species tree analyses. A. Species tree for *Storeria* estimated with SVDquartets from a ddRADseq dataset of unlinked SNPs assembled with a clustering threshold of 90%. Numbers indicate bootstrap support values. B. Species tree for *Storeria* estimated with BPP. Numbers on nodes represent posterior probabilities from the BPP A01 analyses.

The species tree inferred with BPP (Fig. 4B) is congruent with the topology recovered by SVDquartets. However, the relationship between the south-central U.S.A. samples and those from the CCB was strongly supported, with a posterior probability of 1. Again, we recognize that it is potentially problematic to treat the samples from south-central U.S.A. as a clade.

### 3.4. Isolation-by-distance (IBD) and species delimitation

The IBD analysis suggests that *S. dekayi* represents more than one distinct species. The genetic distances estimated within and among samples assigned to different groups from various geographic locations are discontinuous, with genetic differentiation being largely independent of geographic separation (Fig. 5).

**Figure 5.**
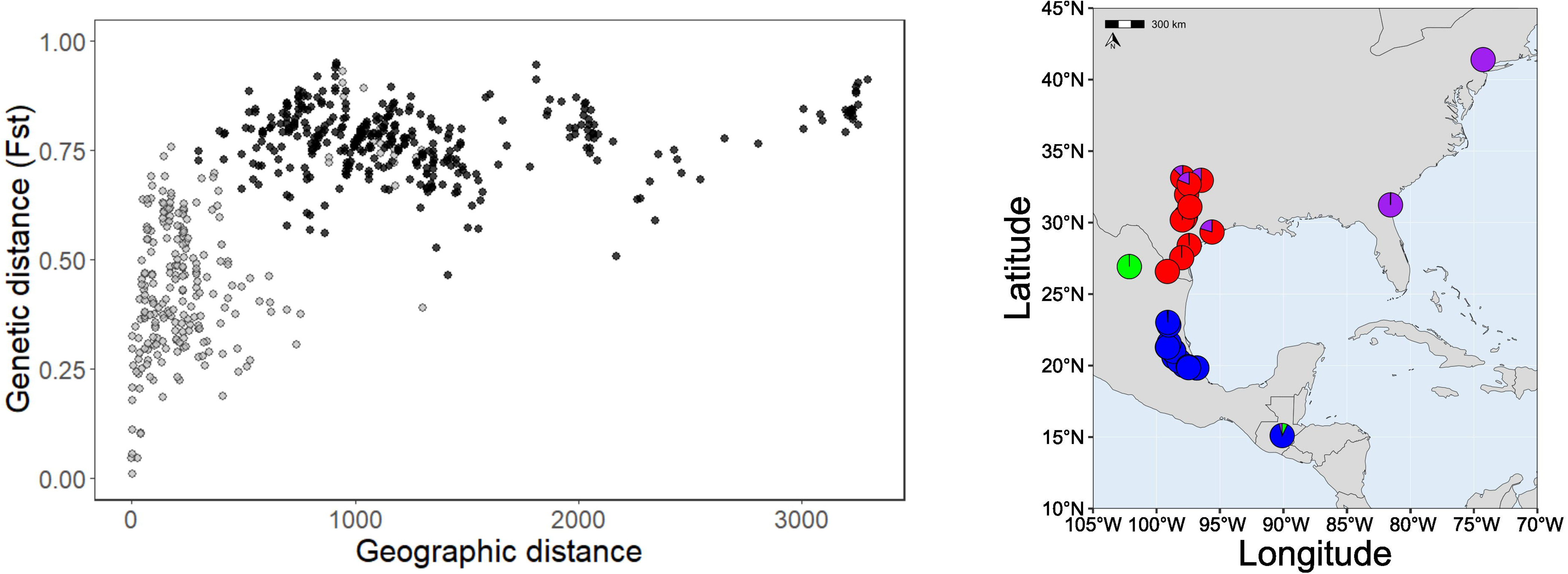
Pairwise genetic distance (FST) between individuals assigned to the established populations, with gray representing FST within populations and black representing FST between different populations, as a function of the geographic distances among them. The relationship between genetic and geographic distances within and among groups is expected to be discontinuous in the presence of distinct species.

The gdi values calculated from the parameters estimated by BPP (Fig. 6A), excluding admixed samples, suggest that the lineages from CCB and *S. dekayi* from Mexico and Central America each represent a distinct species (gdi = 0.997 and 0.873, respectively), whereas the eastern and south-central U.S.A. lineages fall within the zone of ambiguity (gdi = 0.338 and 0.265, respectively). In contrast, the estimates including admixed samples (Fig. 6B) indicate that only CCB represents an independent evolutionary lineage (gdi = 0.871), whereas the *S. dekayi* lineages from Mexico and Central America and from the eastern U.S.A. fall within the zone of ambiguity (gdi = 0.673 and 0.296, respectively). The south-central U.S.A. lineage falls below the 0.2 threshold (gdi = 0.068).

**Figure 6.**
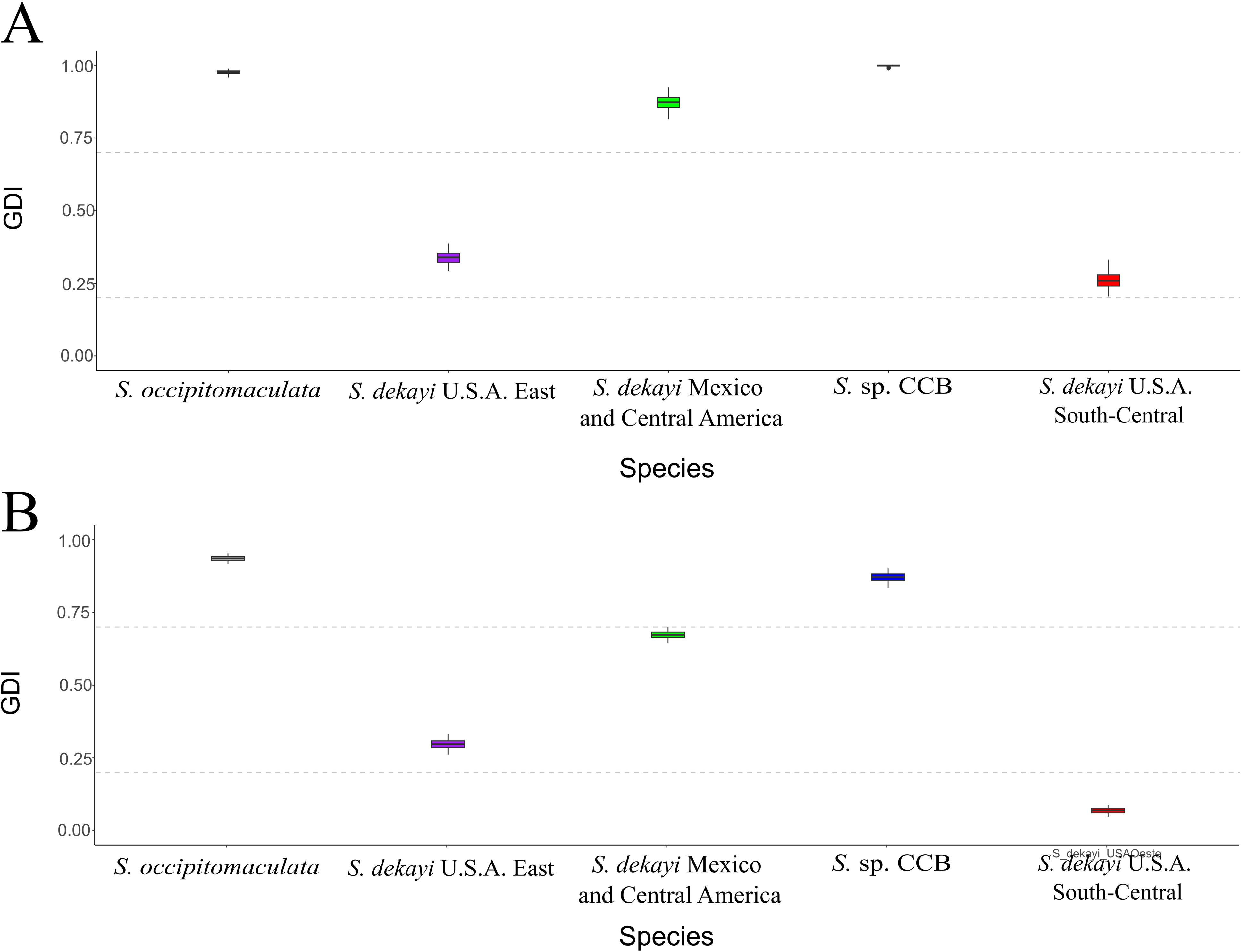
Heuristic genealogical divergence index (gdi) values for *Storeria occipitomaculata* and putative species within *S. dekayi*. The gdis were calculated using the estimates for population sizes (theta) and divergence times (tau) from the analysis of 500 loci in BPP. The tree from a BPP 01 analysis was used as guide tree. Gdi values between gdi = 0.2 and gdi = 0.7 (dotted lines) indicate that species delimitation in this interval is ambiguous. A. gdi values excluding highly admixed samples. B. gdi values including highly admixed samples.

Starting from a five-population species tree (*S. occipitomaculata*, eastern U.S.A., south-central U.S.A., Mexico and Central America, and CCB), and excluding the highly admixed samples, the hhsd approach supported the recognition of five species, with a high migration rate (M = 0.190; Table S5) observed between the eastern U.S.A. and south-central U.S.A. lineages. In contrast, the split algorithm suggested that the entire *S. dekayi* complex should be considered a single species, with a high migration rate (M = 0.106; Table S6) between eastern U.S.A. and the combined group of populations from Mexico and Central America, CCB, and south-central U.S.A.

On the other hand, when including the highly admixed samples and starting from the same five-population species tree (*S. occipitomaculata*, eastern U.S.A., south-central U.S.A., Mexico and Central America, and CCB), the merge algorithm of hhsd recognized two distinct lineages within the *S. dekayi* complex: the eastern U.S.A. lineage and a combined group from Mexico and Central America, CCB, and south-central U.S.A. The split algorithm recovered the same lineages, and both algorithms showed high migration rates (merge M = 0.174; split M = 0.175; Tables S7 and S8. respectively) from the eastern U.S.A. populations to the group composed of Mexico and Central America, CCB, and south-central U.S.A.

### 3.5. Hybridization

Phylonetworks analyses revealed that all models involving reticulation events (h > 0) fit the data better than models based on strictly bifurcating trees (h = 0). Negative pseudolikelihood scores decreased from h = 0 to h = 1 and remained essentially unchanged, or showed only minimal improvement, for h > 1 (Fig. S6), suggesting that the best-fitting phylogenetic model includes a single introgression event. This network indicates an introgression event from *S. dekayi* from south-central U.S.A. into those from eastern U.S.A., with an inheritance probability of 33.3% (γ = 0.333; Fig. 7).

**Figure 7.**
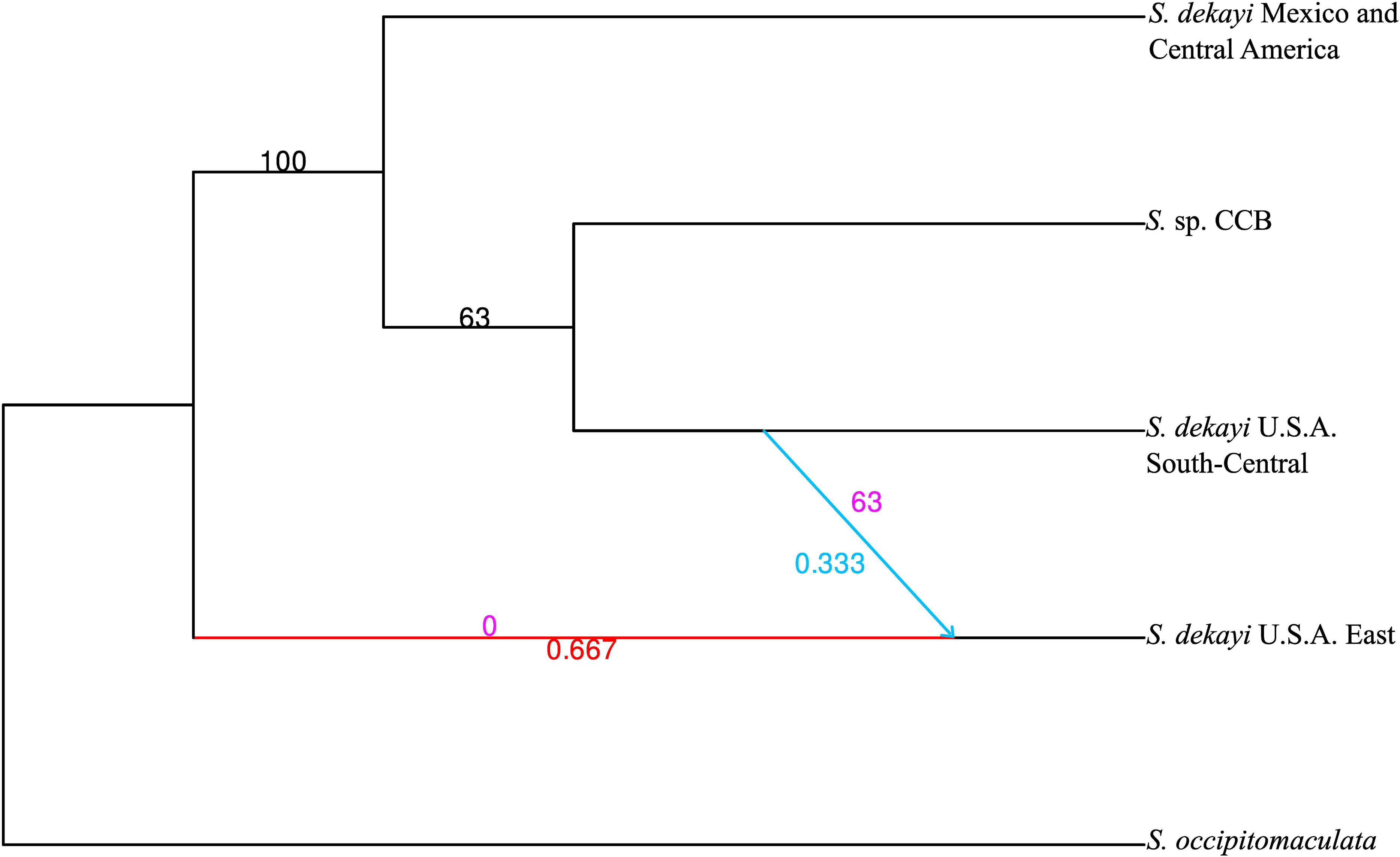
Phylogenetic network inferred by PhyloNetworks with hmax=1. Bootstrap values are based on 100 bootstrap replicates. The blue arrow indicates an inferred hybridization event and its direction. Red and blue numbers are the estimated proportions of genes (heritabilities) contributed by the major and minor hybrid edges, respectively, to the hybrid taxon. Black numbers are bootstrap support values for the non-hybrid nodes, and purple numbers are bootstrap support values for the major and minor parent edges of the hybrid taxon.

### 3.6. Divergence-time estimation

The split between *S*. *occipitomaculata* and the *S*. *dekayi* complex (Fig. 8; Table 2) was estimated at approximately 21.67 Mya (Node A; 95% HPD = 15.35–28.24). Within the *S*. *dekayi* complex, the earliest divergence occurred around 5.46 Mya (Node B; 95% HPD = 3.91–7.11), separating the eastern U.S.A. lineage from the clade comprising Mexico and Central America, south-central U.S.A. and *S*. sp. CCB. Subsequently, the divergence between the Mexican and Central American populations and the south-central U.S.A. and *S*. sp. CCB populations was estimated at 2.69 Mya (Node C; 95% HPD = 1.84–3.58). Finally, the split between the south-central U.S.A. populations of *S*. *dekayi* and *S*. sp. CCB was estimated at 2.40 Mya (Node D; 95% HPD = 1.64–3.24).

**Figure 8.**
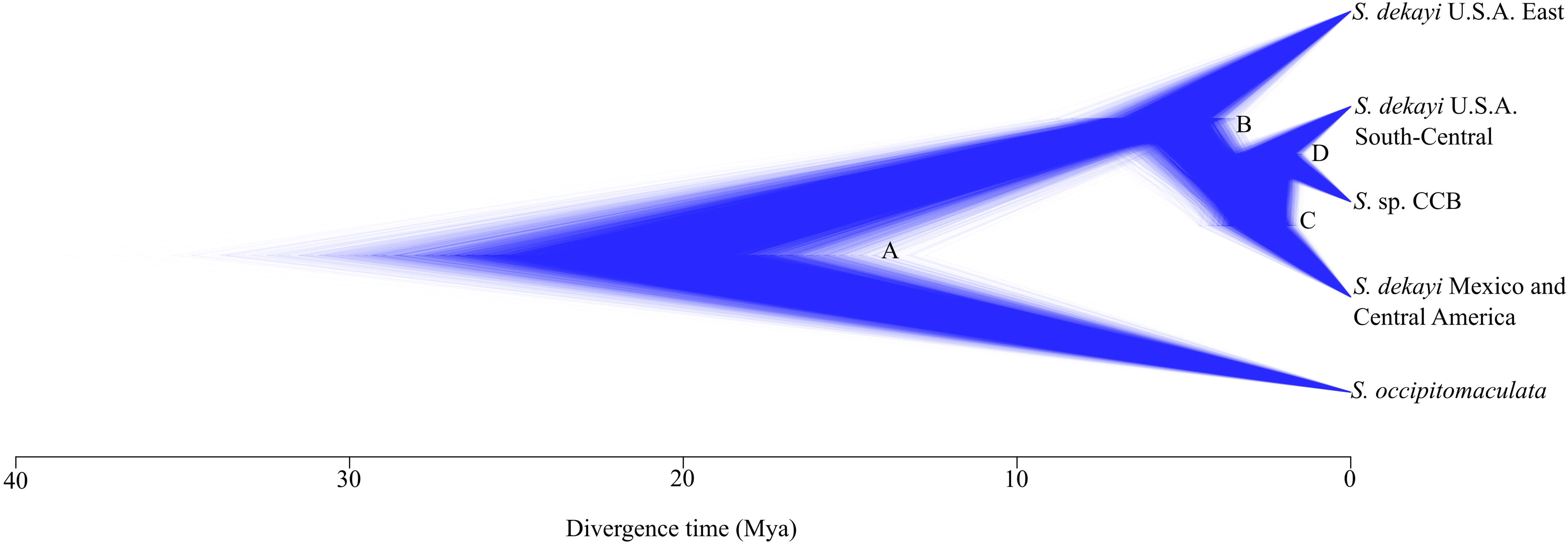
Divergence time estimates from Multi Species Coalescent A00 analysis in BPP. Node labels correspond to those in Table 4. Node A: divergence between *S*. *occipitomaculata* and the *S*. *dekayi* complex. Node B: divergence between *S*. *dekayi* from eastern U.S.A. and *S. dekayi* from Mexico and Central America *+ S.* sp. CCB + *S. dekayi* from south-central U.S.A. Node C: divergence between *S. dekayi* from Mexico and Central America and *S.* sp. CCB + *S. dekayi* from south-central U.S.A. Node D: divergence time between *S.* sp. CCB and *S. dekayi* from south-central U.S.A.

**Table 2.** Estimated divergence times using BPP. Node labels correspond to those in. **Fig. 8**.

| Node | Estimated Age (Mya) | HDP-95% |
| --- | --- | --- |
| A | 21.67 | 15.35–28.24 |
| B | 5.46 | 3.91–7.11 |
| C | 2.69 | 1.84–3.58 |
| D | 2.40 | 1.64–3.24 |

### 3.7. Morphological comparisons

Biplots of the first three principal components (PCs; Fig. 9) explained 70.83% of the total variation in the morphological characters (Table S9): the first (53.14% of variation) mainly represented variation in head length, mandible length, and maximum parietal scale length; the second (9.99% of variation) represented variation in the number of ventral and subcaudal scales and vertical eye diameter; and the third (7.69% of variation) was represented by the number of subcaudal scales and total length. The populations from CCB were fully separated from those in eastern and south-central U.S.A. along PC2 (Fig. 9A), but all three sets of populations were largely overlapping with those from Mexico and Central America on this axis. On PC1, there was both some separation and some overlap between populations from CCB, eastern U.S.A., and south-central U.S.A. and those from Mexico and Central America (Fig. 9A, B), Populations were largely overlapping on PC3 (Fig. 9B, C).

**Figure 9.**
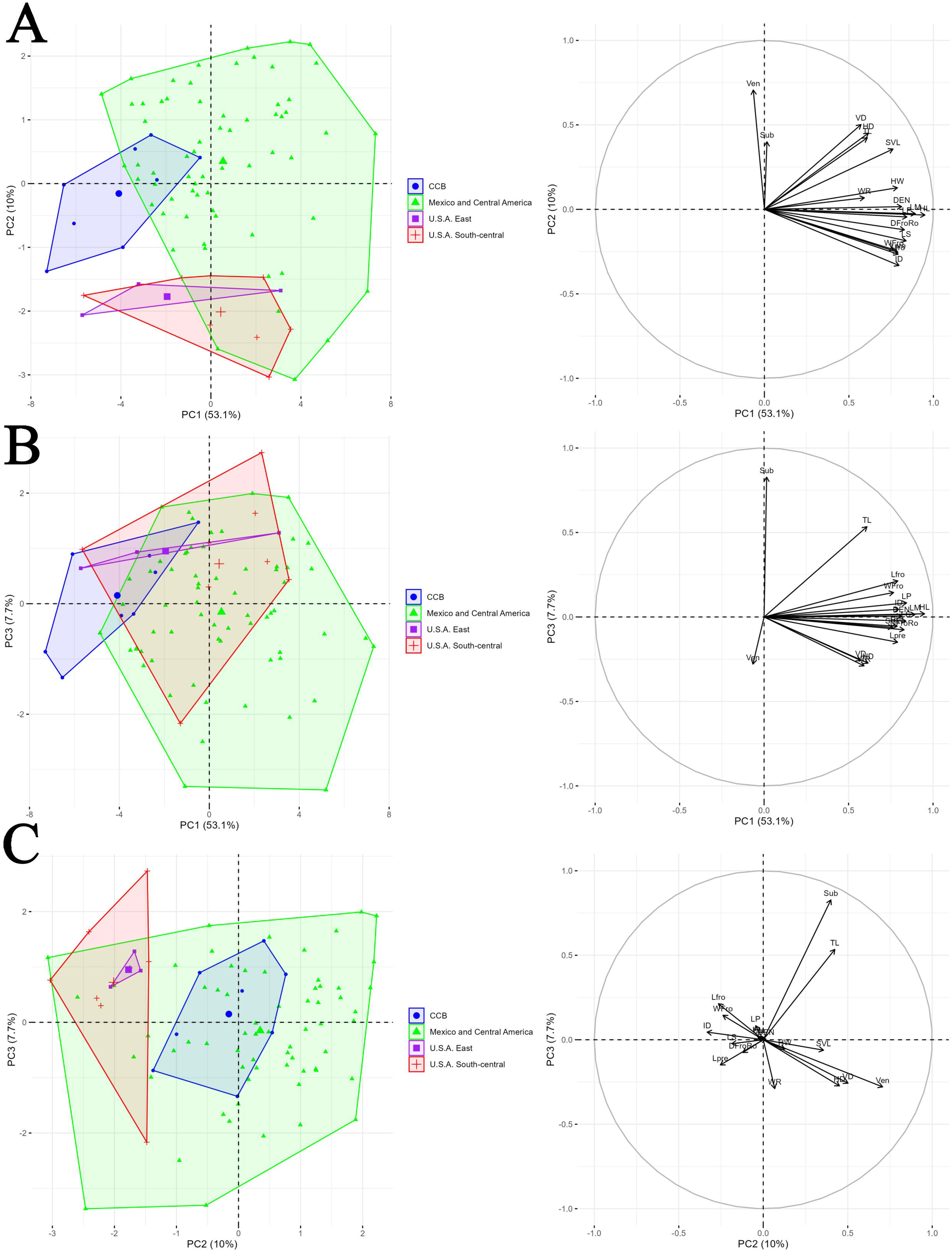
Results of PCA of morphological data. A) Biplot visualization of PC axes 1 and 2 and variable factor map illustrating variable loadings for PC axes 1 and 2. B) Biplot visualization of PC axes 1 and 3 and variable factor map illustrating variable loadings for PC axes 1 and 3. C) Biplot visualization of PC axes 2 and 3 and variable factor map illustrating variable loadings for PC axes 2 and 3.

## 4. Discussion

Our analyses of RADseq and morphological data suggest that the *S. dekayi* complex consists of multiple lineages, but the number of lineages that are distinct varies across analyses. Our STRUCTURE and PCA analyses of the RADseq data (Fig. 2) suggest that populations from Mexico and Central America are distinct from all others. Phylogenetic analyses of the concatenated RADseq data show that populations from Mexico and Central America form a strongly supported clade that is sister to the populations from the Cuatro Cienegas Basin (CCB), but that this clade is nested among populations from the eastern and south-central U.S.A. (Fig. 3). Species-tree analyses show the CCB population as sister to those from south-central U.S.A. (Fig. 4), with populations from Mexico and Central America and eastern U.S.A. as successive outgroups. Analyses using the gdi approach suggest that populations from CCB and those from Mexico and Central America are distinct species (Fig. 6). Morphological analyses show that populations from CCB and those from eastern and south-central U.S.A. are distinct from each other, but not from those in Mexico and Central America (Fig. 9). Overall, given the conflicts among these results, we tentatively recognize only two species: the CCB lineage and *Storeria dekayi*, which includes all remaining populations from the U.S.A., Mexico, and Central America.

### 4.1. Phylogenetic relationships, population structure and gene flow

The phylogenetic relationships among *Storeria* species recovered in this study are consistent with previous species-level studies (Pyron et al., 2016; Nuñez et al., 2023), which identified *S. dekayi* as the sister group to *S. victa*. However, our focus here was on relationships within *S. dekayi*, especially those populations from Mexico and Central America. Our ddRADseq-based analyses revealed that *S. dekayi* is paraphyletic with respect to *S.* sp. CCB, which is congruent with previous findings from mitochondrial data (García-Vazquez, 2020; Vite-Hernandez, 2023). Similar patterns involving endemic taxa from Cuatro Cienegas have been observed in other reptiles (e.g., *Terrapene coahuila*, *Gerrhonotus mccoyi* and *Scincella kikaapoa*), suggesting historical isolation followed by secondary contact (Scheinvar et al., 2020; Blair et al., 2022) or paraphyly of the ancestors of the CCB endemics (e.g., *S. dekayi*) due to incomplete lineage sorting.

The STRUCTURE and PCA analyses of the RADseq data revealed three main clusters in the *S. dekayi* complex (Fig. 2): one from the eastern U.S.A., another from south-central U.S.A., and a third from Mexico and Central America. However, there was clearly admixture between populations from eastern and south-central U.S.A., based on the PCA and STRUCTURE analyses (with K=3, 4, and 5) Subsequently, four groups were used as hypotheses in the species-tree analyses given that the CCB population was differentiated in the ML analyses and in STRUCTURE analyses with K=4 and 5. In the resulting species-tree estimates, the south-central U.S.A. samples were placed as the sister group to the CCB species (Fig. 4). By contrast, in the concatenated ML tree (Fig. 3B), the eastern U.S.A. samples were strongly supported as the sister taxon to the other *S. dekayi* populations, the south-central U.S.A. populations were strongly supported as being paraphyletic with respect to the Mexican populations, and the population from CCB was strongly supported as sister to the populations from eastern Mexico and Central America.

The discrepancy between the species trees and ML tree may be attributed to limited geographic sampling which can create the appearance of distinct genetic groups (Barley et al., 2018). As Chambers and Hillis (2020) noted, restricted sampling combined with a strong conflicting signal of introgression can lead to an overestimation of the number of distinct lineages by several methods such as BPP or STRUCTURE. In our case, the results may reflect high admixture among genetic groups (as seen in STRUCTURE; Fig. 2A), historical introgression (which can mislead phylogenetic inference; Esquerre et al., 2019), or incomplete lineage sorting. The group formed by the samples from the south-central U.S.A. includes individuals with genetic contributions from the eastern U.S.A. populations, which may be due to a zone of overlap between populations or an area of intergradation that indicates intraspecific variation (Hillis et al., 2021). This may explain the paraphyly of the south-central U.S.A. populations with respect to the clade of CCB and the Mexican and Central American populations in the ML tree (Fig. 3), as any degree of hybridization between species can lead to paraphyly (Hillis, 2022). A broader sampling effort in the contact zones between the eastern and south-central U.S.A. populations would help clarify the relationships between them, whether they represent a case of failed speciation with two populations experiencing widespread gene flow (Schield et al., 2015) or a “genetic sink” that restricts gene flow between species (Yanchukov et al., 2006; Chambers and Hillis, 2020). Moreover, genomic data can reveal evidence of introgression over time (San Jose et al., 2023). Our PhyloNetworks results suggested a hybridization event in which *S. dekayi* populations from the eastern U.S.A. received approximately one-third of their genomes (γ = 0.333) from the populations from the south-central U.S.A. (but note that this is based on only a sampling of the genome from RADseq data). A similar pattern was inferred between the snakes *Lampropeltis gentilis* and *L*. *annulata* (Ruane et al., 2014).

The geographically structured distribution of genetic variation observed in the STRUCTURE analysis is not unexpected, given the large geographic distances between populations. Additionally, the apparent lack of admixture between the lineages from Mexico and Central America and the U.S.A. could reflect a failure to sample intermediate populations in southern Texas, potentially leading to an overestimation of species diversity, as seen in the eastern and south-central U.S.A. populations. Therefore, an IBD analysis was used to determine whether geographic isolation alone is limiting gene flow. The genetic and geographic distances, both within and between lineages, showed a discontinuous pattern, and genetic distance is independent of geographic distance (Fig. 5). This result may indicate mechanisms that restrict gene flow, such as reproductive isolation, supporting species delimitation and aligning with expectations of evolutionary divergence (Prates et al., 2024). We also note that within the Mexico-Central America clade, the sister to all other populations is from Guatemala, not the populations adjacent to Texas in Tamaulipas (Fig. 3B).

### 4.2. Species limits and taxonomy

We found substantial genetic divergence among populations traditionally referred to as *Storeria dekayi*. Historical morphological assessments distinguished Mexican and Central American populations from other populations based on characters such as the presence or absence of a black temporal line, a U-shaped orbital mark, and differences in ventral scale counts (Trapido, 1944; Anderson, 1961). However, Sabath and Sabath (1969) noted the difficulty of identifying *S. dekayi* subspecies due to morphological overlap and potential hybridization, an observation corroborated here by the shared genetic variation and evidence of admixture among lineages. *Storeria* sp. CCB is morphologically distinguished from *S. dekayi* by its smaller size, lower number of ventral and subcaudal scales, and unique coloration pattern (Vite-Hernandez, 2023). As noted by Trapido (1944), there is a tendency for southern populations to have a higher number of ventral scales, which is consistent with the specimens examined here from Mexico and Central America (*S. dekayi* south-central U.S.A. (*n*=8): 123–142, *x̅* = 133.00; *S. dekayi* eastern U.S.A. *(n*=3): 126–137, *x̅* = 133.00; S. sp. CCB (*n*=8): 129–137, *x̅* = 133.25; *S. dekayi* Mexico and Central America (*n*=65): 119–149, *x̅* = 137.49). Nevertheless, further morphological analyses are warranted to identify consistent diagnostic traits among these lineages.

Based on gdi values (Fig. 6), only the CCB lineage was genetically distinct enough to be considered evolutionarily independent (gdi > 0.7), whereas the populations from Mexico and eastern U.S.A. fell within the ambiguity zone. The populations from south-central U.S.A. lineage were below the 0.2 threshold, suggesting that they are not a distinct species (Leache et al., 2017). Leache et al. (2019) identified two limitations of the gdi metric: (1) it may yield discordant species assessments between sister lineages with differing levels of genetic diversity, and (2) its estimates are sensitive to θ values (population sizes), with higher values in lineages exhibiting low intraspecific variation. The CCB lineage, being narrowly distributed, likely achieved genetic distinctiveness more rapidly, whereas the ambiguous (lower) gdi values for the populations from the eastern and south-central U.S.A. may reflect their broader distributions and thus larger effective population size. The paraphyly of the south-central U.S.A. populations with respect to the eastern U.S.A. populations observed in the concatenated ML analysis (Fig. 3B) may be due (in part) to admixture between these lineages. However, the strongly supported paraphyly of the south-central U.S.A. populations may also represent incomplete lineage sorting. It is well known that diverging sister species go through a life cycle in which alleles are initially polyphyletic, then paraphyletic, and then monophyletic, with the speed of these transitions depending on the population size (e.g. Rosenberg, 2003). Thus, it is unsurprising that the widely distributed *S. dekayi* populations in North America remain paraphyletic, whereas the more narrowly distributed lineages in Cuatro Cienegas and in eastern Mexico and Central America have each become monophyletic. We consider it unlikely that the paraphyletic series of lineages in south-central U.S.A. represent additional cryptic lineages, given that the putative cryptic species do not appear to be geographically coherent. The discrepancy between the results given by the *merge* and *split* algorithms in hhsd (Kornai et al., 2024) may be due to the broad uncertainty interval in the *gdi* (0.2 < gdi < 0.7) and the high migration rate between the south-central U.S.A. and eastern U.S.A. populations.

Taken together, our results support the recognition of two or possibly three distinct species. One of these is the undescribed species of *Storeria* endemic to the Cuatro Cienegas Basin. Based on our analyses here, this species was supported as a distinct lineage based on gdi (Fig. 6), STRUCTURE analyses (with K=4 and 5; Fig. 2), and concatenated ML analyses (Fig. 3). It was also morphologically distinct from populations in eastern and south-central U.S.A. based on PCA of the morphological data (Fig. 9). In addition, it exhibits a set of unique characteristics that distinguish it from other Storeria species, including body size and color pattern, as well as its restricted distribution in the wetlands of the CCB region surrounded by desert (Vite-Hernandez, 2023).

*Storeria dekayi* is tentatively considered a single, widely distributed species. We think that there is little evidence to suggest that populations from eastern and south-central U.S.A. represent distinct species, since there appears to be extensive hybridization between them (Figs. 2, 3), given that the south-central populations do not form a monophyletic group (Fig. 3), and given the non-significant gdi results (Fig. 6). On the other hand, there is far more evidence for recognizing the populations from Mexico and Central America as distinct. These populations form a distinct cluster in the STRUCTURE and PCA of the RADseq data (Fig. 2), are strongly supported as monophyletic in the concatenated ML analyses (Fig. 3), and in the gdi analyses are significantly supported as distinct species (Fig. 6A) or are almost significantly supported (Fig. 6B). We did find morphological overlap between the populations from Mexico and Central America and other populations in the morphological PCA (Fig. 9), but this merely indicates that these populations may represent a (semi-) morphologically cryptic species. Overall, we think that there is ample evidence to recognize populations of *S. dekayi* from Mexico and Central America as a distinct species (for which the name *S. tropica*) is available. Although one could argue that more sampling is needed in the unsampled area between the Mexican and U.S. populations in northern Mexico (Fig. 3), it is not clear if *S. dekayi* occurs there.

The apparent paraphyly of *Storeria dekayi* may be due to the fact that our sampling did not include representatives of all described subspecies. A more comprehensive assessment will require broader geographic sampling. Therefore, we tentatively recognize only two species: *S. dekayi* and the distinct lineage from Cuatro Cienegas.

### 4.3. Divergence times

Divergence time estimates from BPP suggest that the separation of populations in eastern and south-central U.S.A. occurred during the Pleistocene, possibly driven by an interglacial eluviation event that expanded the Mississippi River floodplain (Thornbury, 1965, Soltis et al., 2006). The Mississippi River has been recognized as a major phylogeographic barrier in numerous taxa (Lemmon et al., 2007; Burbrink et al., 2008; Pyron et al., 2009). Divergence between the Mexico-Central America lineage and the south-central U.S.A. population may have occurred during the late Pliocene to early Pleistocene, associated with the establishment of the Rio Grande drainage system (Gustavson, 1991).

The split between the CCB and the south-central U.S.A. lineages also dates from the Pleistocene, a period conducive to range shifts and episodic gene flow due to glacial cycles and forest expansions and retractions (Bryson et al., 2011). These climatic fluctuations over the past 2.5 million years played a significant role in shaping species distributions and genetic diversity in North America (Gamez and Castellanos-Morales, 2019; Scheinvar et al., 2020). Populations were reduced and confined to Pleistocene refugia, where genetic drift and local adaptation promoted divergence (Myers et al., 2019). Upon climate amelioration, range expansions from different refugia led to secondary contact, increasing intrapopulation variation and producing complex genetic patterns (Myers et al., 2019). These refugia—documented in many taxa—are recognized as reservoirs of genetic diversity (Morafka, 1977; Metcalf, 1977; Elias et al., 1995; Contreras-Balderas et al., 2007). Cuatro Cienegas is one such refugium and has been identified as a key genetic diversity reservoir for North American biota (Gamez and Castellanos-Morales, 2019).

We note that our estimate for the age of *Storeria* is substantially older than other recent estimates that incorporate broad species sampling and multiple fossil calibration. For example, Zheng and Wiens (2016) estimated that *S. occipitomaculata* and *S. dekayi* diverged 9.11 Mya, and that all North American natricines were only 15 Myr old (crown-group age). Here, we estimated that *S. occipitomaculata* and *S. dekayi* diverged 21.67 Mya (Table 2). Nuñez et al. (2023) estimated the crown-group age of North American natricines at 21.57 Mya, with the crown-group age of *Storeria* <3 Mya. Thus, the ages estimated here should be taken with much caution, because they lack external fossil calibrations.

## Acknowledgements

This paper serves as a fulfillment of VGCS for obtaining a M.Sc. degree in the Posgrado en Ciencias Biólogicas from Universidad Nacional Autónoma de Mexico. We thank the SECIHTI for funding and for the support of this research through a graduate scholarship to VGCS (CVU 1249510). Support for fieldwork was provided by grants from DGAPA, UNAM (PAPIIT-IN 218022 and PAPIME PE208422) to U. O. García-Vazquez, and Richard Glider Graduate School (Theodore Roosevelt Memorial Fund) to VGCS. MTO thanks the support from SECIHTI (Estancias Posdoctorales por Mexico para la Formación y Consolidación de las y los Investigadores por Mexico; CVU131802). For tissue and specimen loans we thank the following institutions: Sam Noble Oklahoma Museum of Natural History, University of Oklahoma, Museo de Zoología FES Zaragoza, UNAM, UTA Amphibian and Reptile Diversity Research Center, American Museum of Natural History, Texas Natural History Collections, Instituto Tecnológico de Huejutla, University of Florida, Florida Museum of Natural History, University of Kansas Biodiversity Institute Herpetology Collection, Museo de Zoología Facultad de Ciencias UNAM. We thank C. Grünwald, A. E. Valdenegro-Brito, J. C. Sanchez-García, A. Garcia-Vinalay for assistance in the field and for donating several samples; and R. G. Martínez-Fuentes, J. C. Sanchez-García, D. I. Sanchez-Aguilar, A. E. Valdenegro-Brito, A. García-Vinalay, N. N. Avila-Jimenez, and A. Vite-Hernandez for their help with laboratory work. All contributors observed state, federal, and international laws concerning the collection and transport of live or preserved specimens. All specimens of Mexico were collected under a collecting permit provided by the Secretaría de Medio Ambiente y Recursos Naturales (permit number FAUT-0243) to U. O. García-Vazquez.

## Author contribution: CRediT

Victor Gabriel Castillo-Sanchez: Conceptualization, Data curation, Formal analysis, Research, Methodology, Writing – original draft, Writing – review & editing. Adrian Nieto-Montes de Oca: Supervision, Validation, Writing – review & editing. John J. Wiens: Funding acquisition, Validation, Writing – review & editing. Marysol Trujano-Ortega: Funding acquisition, Project administration, Writing – review & editing. Uri O. García-Vazquez: Conceptualization, Formal analysis, Funding acquisition, Methodology, Project administration, Resources, Software, Supervision, Validation, Writing – original draft, Writing – review & editing.

## Notes

### Competing Interest Statement

The authors have declared no competing interest.

