## Supplemental files for "Phylogenomics and species delimitation in the brown snake *Storeria dekayi* (Natricidae)"

**Appendix**

Table S1 Description of ddRAD data assemblies constructed for STRUCTURE analyses under a 94% sequence similarity threshold.

| Assembly name | Number of individuals | Number of loci | Number of SNPs | Missing data (%) |
| --- | --- | --- | --- | --- |
| dekayi94_85min | 40 | 1,163 | 6,537 | 12.57 |
| dekayi94_875min | 40 | 817 | 4,544 | 10.89 |
| dekayi94_90min | 40 | 522 | 2,966 | 9.15 |
| dekayi94_925min | 40 | 317 | 1,805 | 7.51 |
| dekayi94_95min | 40 | 145 | 779 | 5.35 |

Table S2. Description of ddRAD data assemblies constructed for genomic analyses under a 92% sequence similarity threshold.

| Assembly name | Number of individuals | Number of loci | Number of SNPs | Missing data (%) |
| --- | --- | --- | --- | --- |
| dekayi92_20min | 50 | 14,916 | 123,552 | 54.93 |
| dekayi92_30min | 50 | 11,116 | 99,177 | 47.61 |
| dekayi92_40min | 50 | 8,613 | 80,633 | 42.04 |
| dekayi92_50min | 50 | 6,309 | 61,779 | 36.57 |
| dekayi92_60min | 50 | 4,135 | 42,488 | 30.76 |
| dekayi92_70min | 50 | 2,190 | 23,881 | 24.42 |
| dekayi92_80min | 50 | 783 | 8,832 | 17.50 |

Table S3 Specimens of *S. dekayi* examined for morphology. Institutional abbreviations for museums and collections follow Sabaj (2023), except for MZFZ (Museo de Zoología of the Facultad de Estudios Superiores Zaragoza, Universidad Nacional Autónoma de Mexico). The abbreviations ANMO, AEVB, ENS, and CIG refer to field numbers for specimens to be catalogued in the MZFZ.

| ID | Species | Locality | Latitude | Longitude |
| --- | --- | --- | --- | --- |
| AEVB 020 | *S. dekayi* | Mexico: Puebla: Ahuacatlan-Tepango Highway | 20.0107 | -97.8302 |
| ANMO 1442 | *S.* sp. CCB | Mexico: Coahuila: Cuatro Cienegas | 26.9189 | -102.1362 |
| ANMO 4322 | *S. dekayi* | Mexico: San Luis Potosí: 1 km S of Huichihuayan | 21.4781 | -98.9614 |
| ANMO 4345 | *S. dekayi* | Mexico: San Luis Potosí: 4 km N of La Pimienta | 21.5644 | -99.0049 |
| ANMO 4380 | *S. dekayi* | Mexico: Puebla: Zaragoza | 19.7829 | -97.5215 |
| IBH 104 | *S. dekayi* | Mexico: San Luis Potosí: Valles | 22.0146 | -99.0060 |
| IBH 922 | *S. dekayi* | Mexico: Veracruz: Ingenio de el Higo | 21.7805 | -98.4550 |
| IBH 15164 | *S. dekayi* | Mexico: Tamaulipas: Matamoros | 25.6574 | -97.5260 |
| IBH 1569 | *S. dekayi* | United States: Pennsylvania: Warren Co. Linestone Twp. along U. S. Rt. 62, miles N. of Ticlioute | 36.3830 | -93.5850 |
| IBH 19734 | *S. dekayi* | Mexico: Veracruz: Coscomatepec | 19.07109 | -97.0450 |
| IBH 2303 | *S. dekayi* | Mexico: Puebla: Rancho Las Margaritas | 19.3499 | -98.5840 |
| IBH 31445 | *S. dekayi* | Mexico: Veracruz: Tlacuilolan | 19.4150 | -97.0506 |
| IBH 7492 | *S. dekayi* | Mexico: Tamaulipas: Ejido Conrado Castillo | 23.5114 | -99.3470 |
| ITAH 329 | *S. dekayi* | Mexico: Hidalgo: Huejutla | 21.1602 | -98.4164 |
| ITAH 105 | *S. dekayi* | Mexico: Hidalgo: Huejutla | 21.0306 | -98.6682 |
| ITAH 1369 | *S. dekayi* | Mexico: Hidalgo: 5km S of La cabaña, Tlanchinol | 20.9798 | -98.6650 |
| ITAH 1384 | *S. dekayi* | Mexico |  |  |
| ITAH 1389 | *S. dekayi* | Mexico |  |  |
| ITAH 168 | *S. dekayi* | Mexico: Hidalgo: Tlanchinol | 21.0306 | -98.6682 |
| ITAH 171 | *S. dekayi* | Mexico: Hidalgo: Huejutla | 21.1652 | -98.4189 |
| ITAH 172 | *S. dekayi* | Mexico: Hidalgo: Jalcotan | 21.1671 | -98.5393 |
| ITAH 233 | *S. dekayi* | Mexico |  |  |
| ITAH 414 | *S. dekayi* | Mexico: Hidalgo: Huejutla | 21.1392 | -98.4098 |
| ITAH 610 | *S. dekayi* | Mexico: Hidalgo: Atlapexco | 21.0476 | -98.3446 |
| ITAH 934 | *S. dekayi* | Mexico: Hidalgo: Atlapexco | 21.0476 | -98.3446 |
| MZFC 29641 | *S.* sp. CCB | Mexico: Coahuila: Cuatro Ciénegas | 26.9189 | -102.1362 |
| MZFC 29643 | *S.* sp. CCB | Mexico: Coahuila: Cuatro Ciénegas | 26.9189 | -102.1362 |
| MZFC 29644 | *S.* sp. CCB | Mexico: Coahuila: Cuatro Ciénegas | 26.9189 | -102.1362 |
| MZFC 29646 | *S*. sp. CCB | Mexico: Coahuila: Cuatro Ciénegas | 26.9189 | -102.1362 |
| MZFC 29647 | *S*. sp. CCB | Mexico: Coahuila: Cuatro Ciénegas | 26.9189 | -102.1362 |
| MZFC 29650 | *S*. sp. CCB | Mexico: Coahuila: Cuatro Ciénegas | 26.9189 | -102.1362 |
| MZFC 29651 | *S*. sp. CCB | Mexico: Coahuila: Cuatro Ciénegas | 26.9189 | -102.1362 |
| MZFC 14226 | *S. dekayi* | Mexico: Hidalgo: Tenango de Noria | 20.3492 | -98.2062 |
| MZFC 14227 | *S. dekayi* | Mexico: Hidalgo: Tenango de Noria | 20.3492 | -98.2062 |
| MZFC 19275 | *S. dekayi* | Mexico: Querétaro | 21.2955 | -99.1192 |
| MZFC 21019 | *S. dekayi* | Mexico: Hidalgo: Tlanchinol | 21.0608 | -98.6311 |
| MZFC 21020 | *S. dekayi* | Mexico: Hidalgo: Tlanchinol | 21.0608 | -98.6311 |
| MZFC 21021 | *S. dekayi* | Mexico: Hidalgo: Tlanchinol | 21.0608 | -98.6311 |
| MZFC 21022 | *S. dekayi* | Mexico: Hidalgo |  |  |
| MZFC 21258 | *S. dekayi* | Mexico: Hidalgo |  |  |
| UANL 2049 | *S. dekayi* | Mexico |  |  |
| UANL 2490 | *S. dekayi* | Mexico: Nuevo León: Col. Nogalar | 25.7229 | -100.2880 |
| UANL 4119 | *S. dekayi* | Mexico: Tamaulipas: Tula-Ocampo road | 22.9184 | -99.5636 |
| UANL 4138 | *S. dekayi* | Mexico: Tamaulipas: Aldama-Soto la Marina road | 23.4055 | -98.0276 |
| UANL 4455 | *S. dekayi* | Mexico: Tamaulipas: 80 Antiguos Morelos- Nuevo Morelos road | 22.5542 | -99.0919 |
| UANL 6127 | *S. dekayi* | Mexico: Nuevo León: Aldama- Soto la Marina road | 26.1640 | -98.6483 |
| UANL 6417 | *S. dekayi* | Mexico: Tamaulipas: Soto la Marina road | 23.7045 | -98.2070 |
| UANL 6441 | *S. dekayi* | Mexico: Tamaulipas: Soto la Marina-Casas road | 23.7102 | -98.2076 |
| UANL 6442 | *S. dekayi* | Mexico: Tamaulipas: Soto la Marina-Casas road | 23.7102 | -98.2076 |
| UANL 6449 | *S. dekayi* | Mexico: Tamaulipas: Mezquital, km 26 S of Matamoros | 25.6181 | -97.5040 |
| UANL 6558 | *S. dekayi* | Mexico: Tamaulipas: La Pesca | 27.4235 | -99.5687 |
| UANL 6587 | *S. dekayi* | Mexico: Tamaulipas: Arcabuz | 27.4235 | -99.5687 |
| UTACV 13071 | *S. dekayi* | United States: New York: Albany | 42.6511 | -73.7528 |
| UTACV 18307 | *S. dekayi* | United States: South Carolina: Edgefield | 33.7896 | -81.9309 |
| UTACV 21770 | *S. dekayi* | Guatemala: Baja Verapaz: Vicinity La Unión Barrios | 15.1775 | -90.2054 |
| UTACV 21771 | *S. dekayi* | Guatemala: Baja Verapaz: Vicinity La Unión Barrios | 15.1775 | -90.2054 |
| UTACV 26585 | *S. dekayi* | Guatemala: Baja Verapaz | 15.1746 | -90.3708 |
| UTACV 28526 | *S. dekayi* | Guatemala: Baja Verapaz: Vicinity La Unión Barrios | 15.1775 | -90.2054 |
| UTACV 28527 | *S. dekayi* | Guatemala: Baja Verapaz: Niño Perdido | 15.1362 | -90.1809 |
| UTACV 28528 | *S. dekayi* | Guatemala: Baja Verapaz: Niño Perdido | 15.1362 | -90.1809 |
| UTACV 32634 | *S. dekayi* | United States: Texas: Titus | 33.1816 | -95.1022 |
| UTACV 32686 | *S. dekayi* | United States: Texas: Titus | 33.1816 | -95.1022 |
| UTACV 32687 | *S. dekayi* | United States: Texas: Titus | 33.1664 | -94.9721 |
| UTACV 33084 | *S. tropica* | Guatemala: Baja Verapaz: Vicinity La Unión Barrios | 15.1775 | -90.2054 |
| UTACV 33819 | *S. dekayi* | United States: Arkansas | 34.6937 | -94.4541 |
| UTACV 33925 | *S. dekayi* | United States: Texas: Young | 31.8880 | -96.0626 |
| UTACV 34113 | *S. dekayi* | United States: Illinois | 41.8306 | -88.0104 |
| UTACV 35994 | *S. dekayi* | United States: Texas: Milam | 31.4630 | -93.8195 |
| UTACV 38249 | *S. dekayi* | Guatemala: Baja Verapaz: Vicinity La Unión Barrios | 15.1775 | -90.2054 |
| UTACV 54319 | *S. dekayi* | Mexico: San Luis Potosí | 21.4956 | -98.9929 |
| UTACV 7074 | *S. dekayi* | Guatemala: Baja Verapaz: Cerro Quisis | 15.1800 | -90.2205 |
| UTACV 7076 | *S. dekayi* | Guatemala: Baja Verapaz: Cerro Quisis | 15.1800 | -90.2205 |
| UTACV 7080 | *S. dekayi* | Guatemala: Baja Verapaz: Cerro Verde | 15.1716 | -90.2002 |
| UTACV 7743 | *S. dekayi* | Guatemala: Baja Verapaz: Cerro Quisis | 15.1800 | -90.2205 |
| UTACV 7744 | *S. dekayi* | Guatemala: Baja Verapaz: Cerro Verde | 15.1716 | -90.2002 |
| UTACV 7745 | *S. dekayi* | Guatemala: Baja Verapaz: Vicinity La Unión Barrios | 15.1775 | -90.2054 |
| UOGV 4767 | *S. dekayi* | Mexico: Hidalgo: Tiaguistnego | 20.7257 | -98.6261 |
| MZFZ 5189 | *S. dekayi* | Mexico: Veracruz: Yecuatla | 19.8322 | -96.8204 |
| MZFZ 5188 | *S. dekayi* | Mexico: Veracruz: Yecuatla | 19.8322 | -96.8204 |
| MZFZ 5190 | *S. dekayi* | Guatemala: Baja Verapaz: Aldea Santa Cruz | 15.1059 | -90.1013 |
| CIG 816 | *S. dekayi* | Mexico: Tamaulipas: El Limón, E of Ocampo | 22.8361 | -99.0263 |
| CIG 817 | *S. dekayi* | Mexico: Tamaulipas: 2 km W of Hwy. 85 on road to Gomez Farías | 23.0367 | -99.1482 |
| CIG 1160 | *S. dekayi* | Mexico: Veracruz | 19.7425 | -96.8186 |
| CIG 1566 | *S. dekayi* | Mexico: Querétaro: El Madroño | 21.2858 | -99.1426 |

Table S4. Descriptions of traits measured and counted for morphological analyses.

| Character type | Character description |
| --- | --- |
| Morphometric characters | Snout-vent length (SVL), measured from the tip of the snout to the anterior margin of the cloaca. |
|  | Tail length (TL), measured from the posterior margin of the cloaca to the tip of the tail. |
|  | Head length (HL), from the tip of the nose to the margin of the parietal scales. |
|  | Head width (HW). |
|  | Distance from the midline between the frontal and rostral scales (DFroRo). |
|  | Length of prefrontal scales along the midline (LPre). |
|  | Length of frontal scale (LFro). |
|  | Width of frontal scale (WFro). |
|  | Maximum length of parietal scale (LP). |
|  | Horizontal diameter of the eye (HD). |
|  | Vertical diameter of the eye (VD) |
|  | Length of supraocular scale (LS). |
|  | Distance from the eye to the nostril (DEN). |
|  | Length of mandible (LM), from the tip of the snout to the end of the last supralabial scale. |
|  | Interorbital distance (ID). |
|  | Width of rostral scale (WR). |
| Meristic characters | Number of ventral scales (Ven). |
|  | Number of subcaudal scales (Sub). |

Table S5. Estimates of migration rates between *S. dekayi* populations from the merge algorithm of hhsd, excluding admixed samples.

| Donor | Recipient | M (95% HPD CI) |
| --- | --- | --- |
| *S. dekayi* U.S.A. East | *S. dekayi* U.S.A. South-Central | 0.190 (0.0, 0.237) |
| *S. dekayi* Mexico and Central America | *S.* sp. CCB | 0.010 (0.0, 0.018) |
| *S. dekayi* Mexico and Central America | *S. dekayi* U.S.A. South-Central | 0.028 (0.0, 0.060) |
| *S.* sp. CCB | *S. dekayi* Mexico and Central America | 0.002 (0.0, 0.009) |
| *S.* sp. CCB | *S. dekayi* U.S.A. South-Central | 0.0008 (0.0, 0.005) |
| *S. dekayi* U.S.A. South-Central | *S. dekayi* U.S.A. East | 0.003 (0.0, 0.017) |
| *S. dekayi* U.S.A. South-Central | *S. dekayi* Mexico and Central America | 0.066 (0.0, 0.084) |
| *S. dekayi* U.S.A. South-Central | *S.* sp. CCB | 0.049 (0.0, 0.065) |

Table S6. Estimates of migration rates between *S. dekayi* populations from the split algorithm of hhsd, excluding admixed samples

| Donor | Recipient | M (95% HPD CI) |
| --- | --- | --- |
| *S. dekayi* U.S.A. East | *S. dekayi* Mexico and Central America *+ S.* sp. CCB + *S. dekayi* U.S.A. South-Central | 0.106 (0.0, 0.131) |
| *S. dekayi* Mexico and Central America *+ S.* sp. CCB + *S. dekayi* U.S.A. South-Central | *S. dekayi* U.S.A. East | 0.002 (0.0, 0.013) |

Table S7. Estimates of migration rates between *S. dekayi* populations from the merge algorithm of hhsd, with admixed samples.

| Donor | Recipient | M (95% HPD CI) |
| --- | --- | --- |
| *S. dekayi* U.S.A. East | *S. dekayi* Mexico and Central America *+ S.* sp. CCB + *S. dekayi* U.S.A. South-Central | 0.174 (0.145, 0.204) |
| *S. dekayi* Mexico and Central America *+ S.* sp. CCB + *S. dekayi* U.S.A. South-Central | *S. dekayi* U.S.A. East | 0.001 (0.0, 0.007) |

Table S8. Estimates of migration rates between *S. dekayi* populations from the split algorithm of hhsd, with admixed samples.

| Donor | Recipient | M (95% HPD CI) |
| --- | --- | --- |
| *S. dekayi* U.S.A. East | *S. dekayi* Mexico and Central America *+ S.* sp. CCB + *S. dekayi* U.S.A. South-Central | 0.175 (0.145, 0.204) |
| *S. dekayi* Mexico and Central America *+ S.* sp. CCB + *S. dekayi* U.S.A. South-Central | *S. dekayi* U.S.A. East | 0.001 (0.0, 0.006) |

Table S9. Eigenvalues, percentage of variance explained, cumulative percentage of variance explained and eigenvectors (loadings) of each morphological trait for each of the principal components. Traits with the heaviest loadings are in bold.

| Eigenvalues | PC1 | PC2 | PC3 | PC4 | PC5 |
| --- | --- | --- | --- | --- | --- |
| Variance | 9.567 | 1.799 | 1.387 | 1.077 | 0.680 |
| % variance | 53.148 | 9.993 | 7.694 | 5.986 | 3.779 |
| Cum. % variance | 53.148 | 63.141 | 70.835 | 76.821 | 80.599 |
| Eigenvectors | PC1 | PC2 | PC3 | PC4 | PC5 |
| SVL | 0.24604133 | 0.26621152 | -0.05255157 | 0.24624963 | -0.19930148 |
| TL | 0.19689449 | **0.31584052** | **0.45429252** | 0.02981611 | -0.01819487 |
| HL | **0.30717746** | -0.02502439 | 0.01609737 | -0.00570768 | -0.16285230 |
| HW | 0.25457346 | 0.09540670 | -0.04690259 | 0.26928406 | 0.26779319 |
| DFroRo | 0.26793787 | -0.09022106 | -0.06438639 | -0.11592825 | -0.23784117 |
| Lpre | 0.25554725 | -0.18944486 | -0.12689815 | -0.10901219 | **-0.30445461** |
| Lfro | 0.25594285 | -0.19790379 | 0.18119172 | 0.06625116 | -0.10158826 |
| WFro | 0.24786065 | -0.17799568 | 0.12296391 | -0.11638493 | 0.27657336 |
| LP | **0.27266899** | -0.03356381 | 0.07089631 | 0.15551303 | -0.28080335 |
| HD | 0.19905504 | **0.33641401** | -0.23205550 | **-0.43071907** | 0.10595239 |
| VD | 0.18484299 | **0.37427380** | -0.21858043 | **-0.48027955** | 0.05793174 |
| LS | 0.27115699 | -0.13873454 | -0.01998967 | -0.02739625 | 0.01652149 |
| DEN | 0.26260172 | 0.01188965 | 0.00370372 | -0.19581768 | 0.15542572 |
| LM | **0.28889817** | -0.01926627 | 0.01255115 | 0.12356479 | -0.06242477 |
| ID | 0.25758836 | -0.24774387 | 0.03837199 | 0.14222955 | 0.08648775 |
| WR | 0.19125100 | 0.05081922 | -0.24582785 | **0.34341008** | **0.64006217** |
| Ven | -0.02132607 | **0.52687298** | -0.23720874 | **0.43133467** | -0.26337029 |
| Sub | 0.00492202 | 0.29845199 | **0.70342188** | -0.06102115 | 0.13174470 |

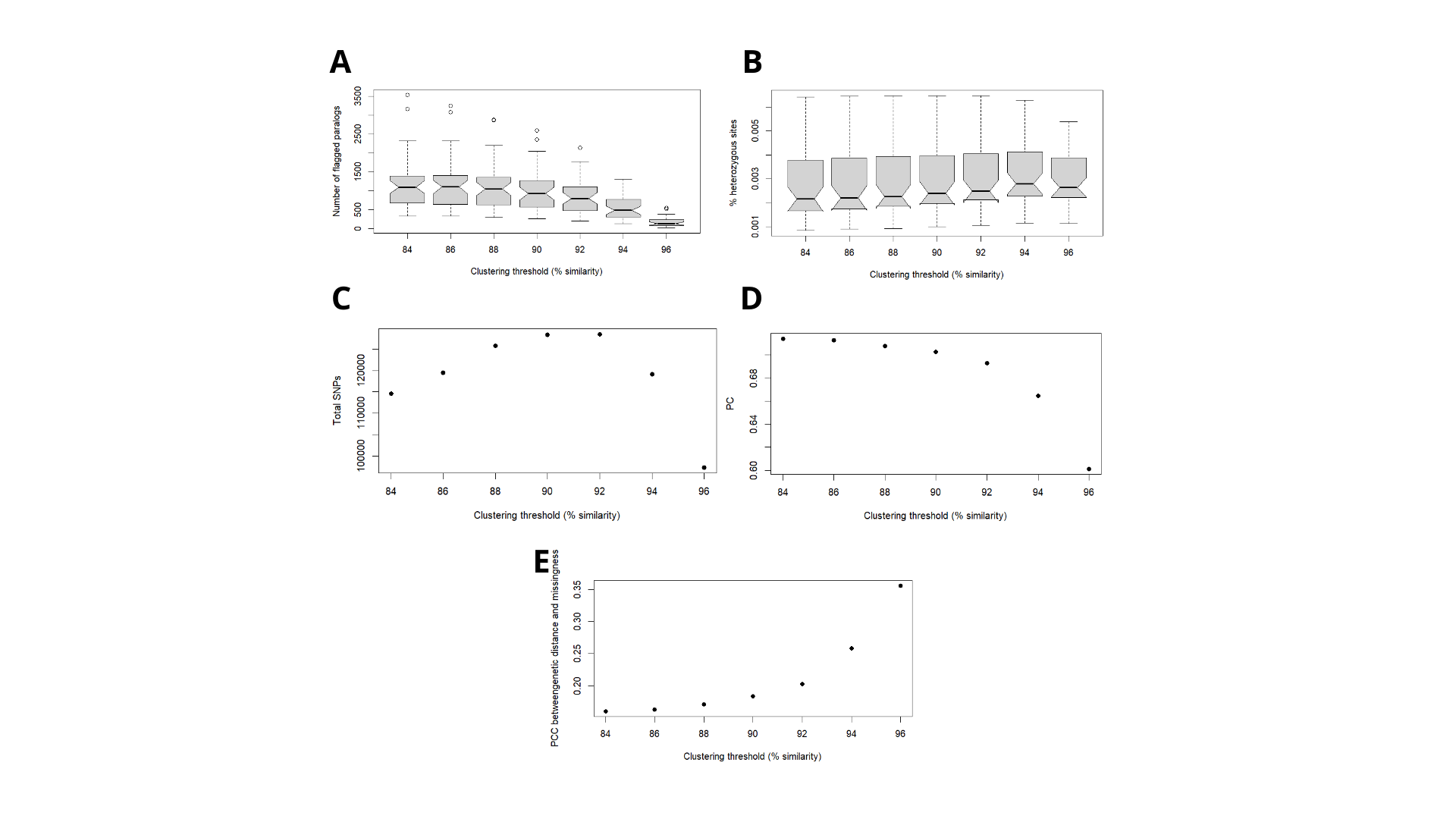

Figure S1. Visualization of metrics calculated for sequence-similarity thresholds ranging from 88% to 96%. (A) Percentage of loci inferred as paralogues, (B) per-individual heterozygosity, (C) total number of SNPs recovered, (D) cumulative variance of biallelic SNPs explained by the first 8 principal components and (E) Pearson's correlation coefficient between pairwise genetic distance and data missingness.

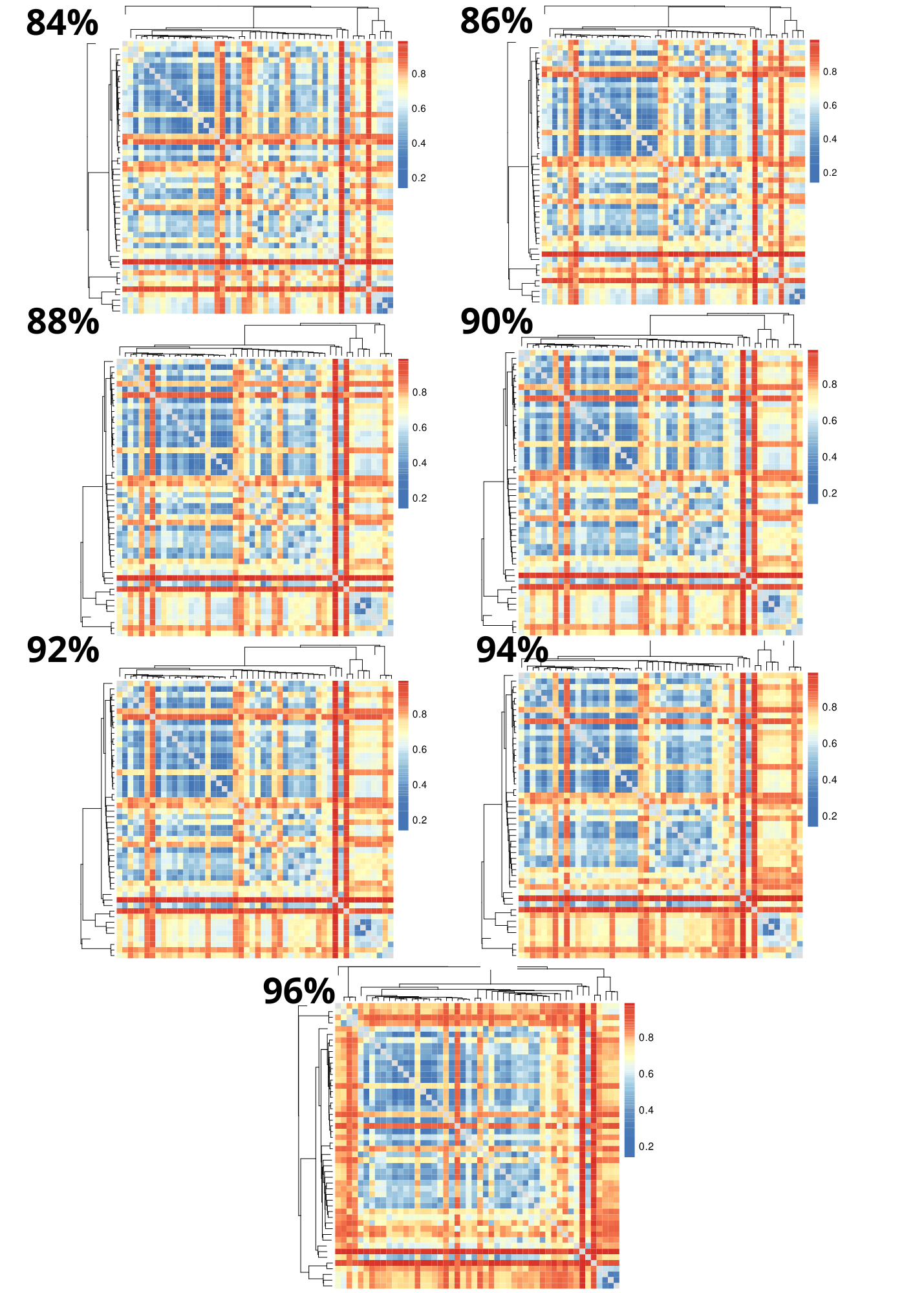

Figure S2. Heat maps visualizing pairwise missing data at clustering thresholds of 88% to 96%. Rows/columns represent individual samples, and dendrograms represent samples clustered by genetic similarity. Colors indicate pairwise fractions of missing data, with cool colors representing low fractions and warm colors representing high fractions.

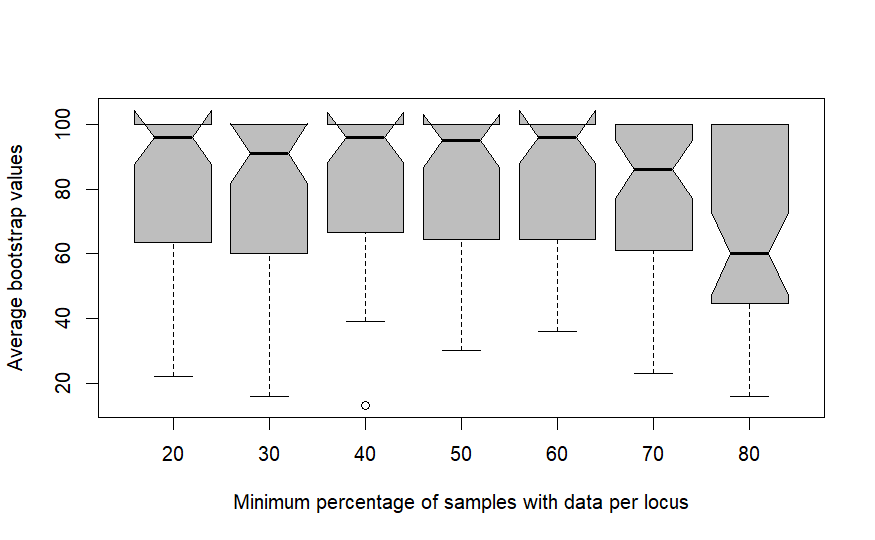

Figure S3. Average bootstrap values for trees inferred from assemblies with varying minimum percentage of samples with data per locus.

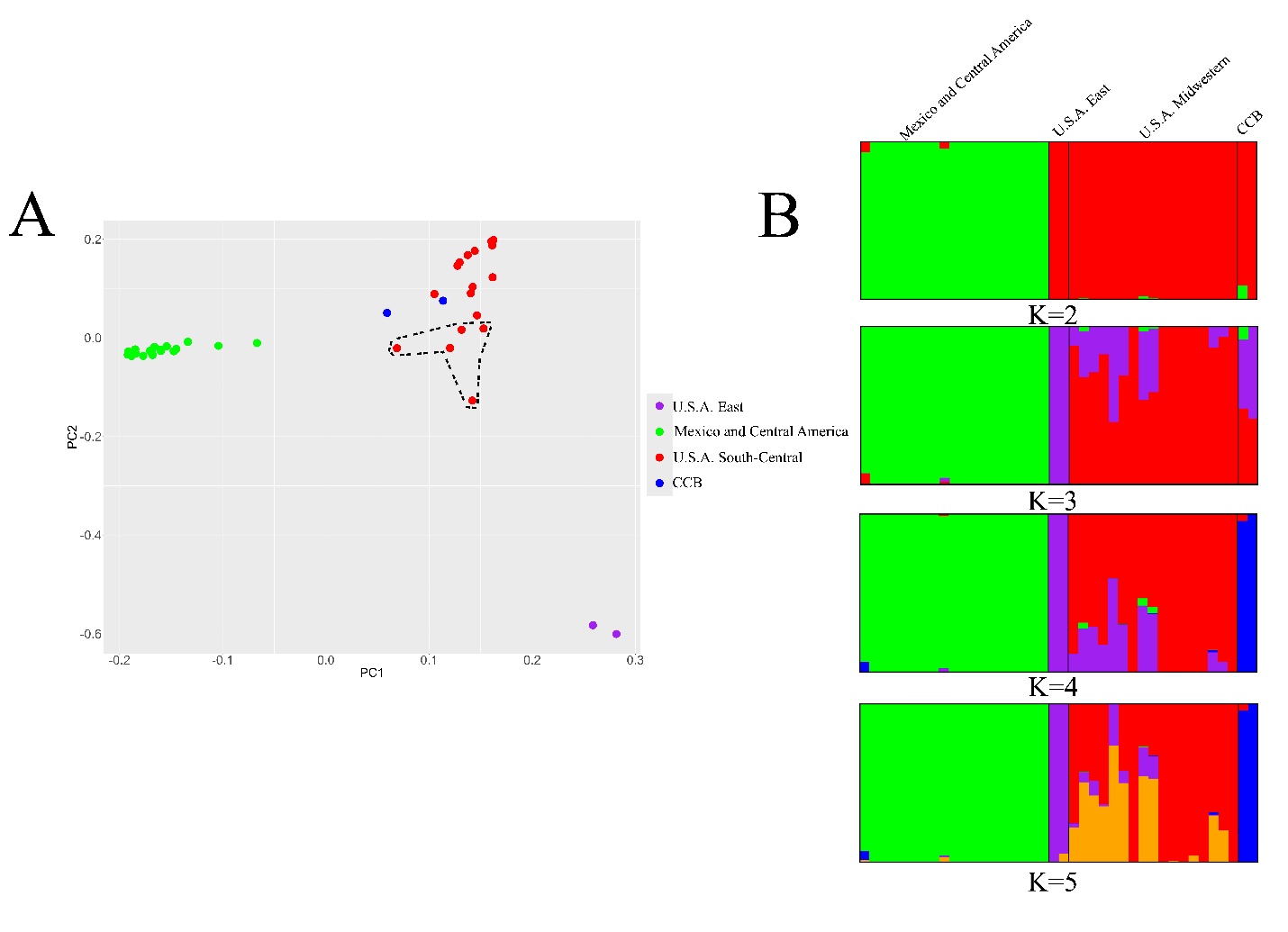

Figure S4. Population structure in *Storeria dekayi* excluding singletons in VCF file A. Population structure inferred using Principal Component Analysis (PCA), with points representing the different individuals colored by cluster. Note the presence of five highly admixed individuals from U.S.A. enclosed by the dashed line. B. Barplot visualization of population structure inferred by STRUCTURE. Each vertical bar represents an individual.

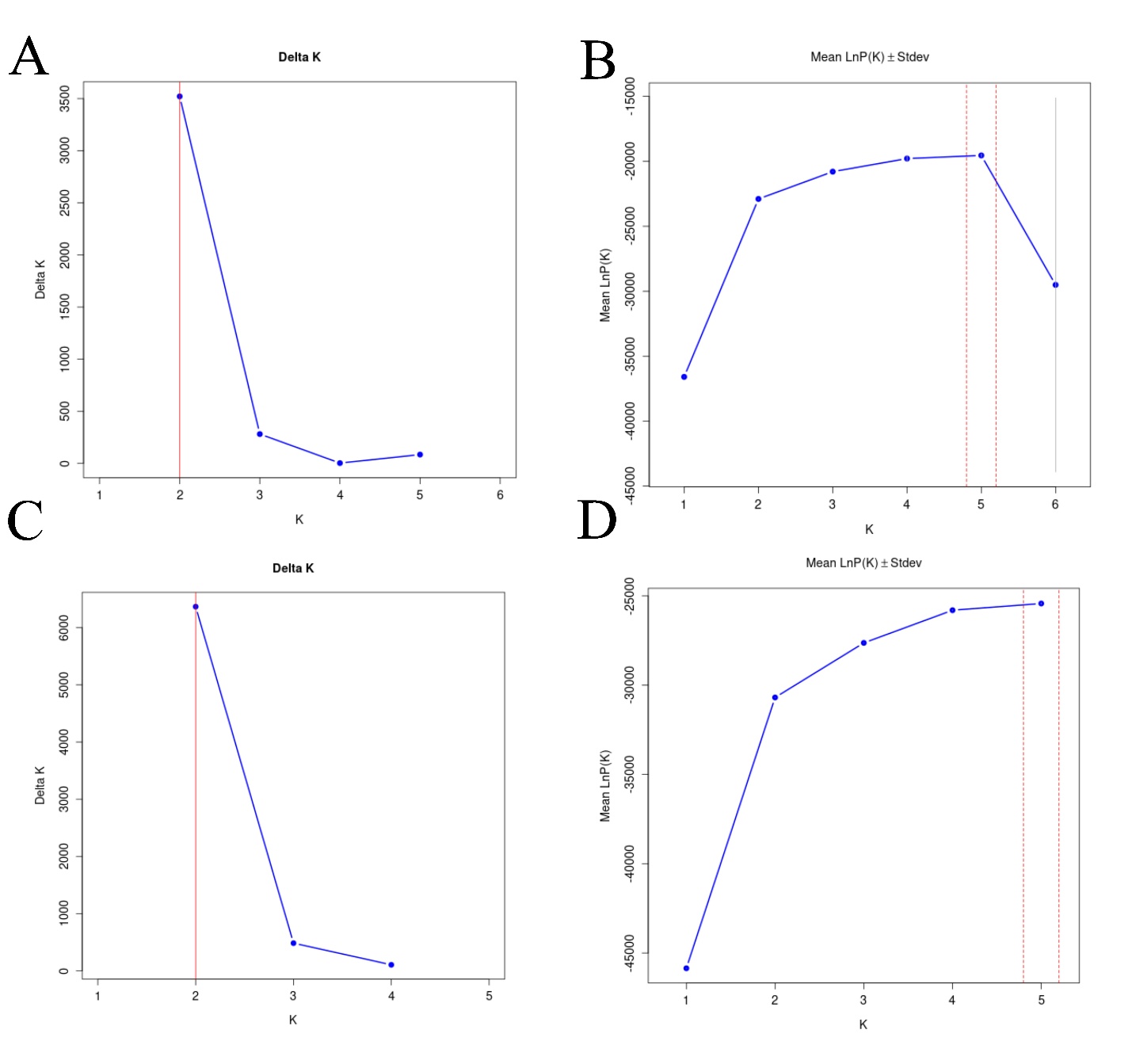

Figure S5. Delta K and ln Pr(D|K) values for varying numbers of K in STRUCTURE analyses of a SNP dataset filtered using different options. A and C: Delta K values in analyses of the SNP dataset filtered with the --maf option and excluding singletons, respectively. B and D: ln Pr(D|K) values in analyses of the SNP dataset filtered with the --maf option and excluding singletons, respectively.

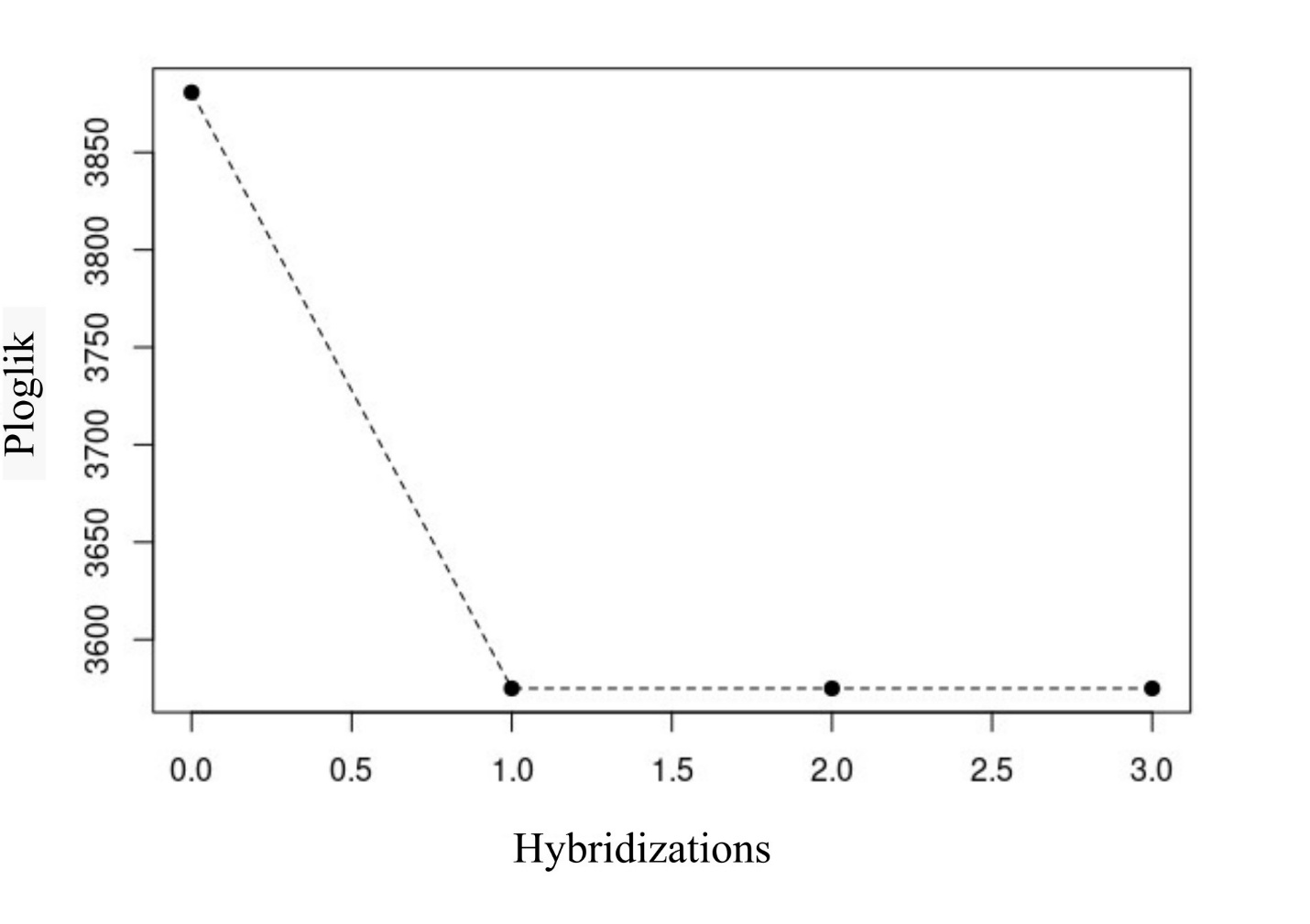

Figure S6. Log-likelihood scores of phylogenetic networks with varying numbers of hybridization events estimated with PhyloNetworks.
